# The phosphate starvation response recruits the TOR pathway to regulate growth in Arabidopsis cell cultures

**DOI:** 10.1101/2021.03.26.437164

**Authors:** Thomas Dobrenel, Sunita Kushwah, Umarah Mubeen, Wouter Jansen, Nicolas Delhomme, Camila Caldana, Johannes Hanson

## Abstract

In eukaryotes, TOR (Target Of Rapamycin) is a conserved regulator of growth that integrates both endogenous and exogenous signals. These signals include the internal nutritional status, and in plants, TOR has been shown to be regulated by carbon, nitrogen and sulfur availability. In this study, we show that in Arabidopsis the TOR pathway also integrates phosphorus availability to actively modulate the cell cycle, which in turn regulates the intracellular content of amino acids and organic acids. We observed a substantial overlap between the phenotypic, metabolic and transcriptomic responses of TOR inactivation and phosphorus starvation in Arabidopsis cell culture. Although phosphorus availability modulates TOR activity, changes in the levels of TOR activity do not alter the expression of marker genes for phosphorus status. These data prompted us to place the sensing of phosphorus availability upstream of the modulation of TOR activity which, in turn, regulates the cell cycle and primary metabolism to adjust plant growth in plants.

## Introduction

The antibiotic rapamycin was described in 1991 as having the capacity to arrest cell division and block cell cycle progression in yeasts (Heitman et al., 1991). Its binding target, conveniently called Target Of Rapamycin (TOR for short), is conserved among all eukaryotic organisms (Dobrenel et al., 2016a) and forms heteromeric protein complexes to exert protein kinase activity resulting in the modulation of various pathways including protein translation, nutrient assimilation, and autophagy, as well as cell division, acting as a master regulator of growth. In plants, the TOR complex includes the proteins TOR, RAPTOR (Regulatory Associated protein of TOR) and LST8 (Lethal with SEC13 8) (Deprost et al., 2005; Moreau et al., 2012).

Strong expression of *TOR* in constantly dividing cells (Menand et al., 2002) and drastic arrest of growth in response to inactivation of the TOR complex components (Deprost et al., 2005, 2007; Menand et al., 2002) supported an early hypothesis of a link between TOR activity and the modulation of cell division in Arabidopsis. More recently, the characterization of a TOR-dependent phosphorylation site in the E2F cell cycle transcription factors and the regulation of *CycB1;1* expression by TOR expression (Li et al., 2017; Xiong et al., 2013), as well as a genetic link between the TOR pathway and the YAK1 kinase resulting in inhibition of expression of the SIAMESE-RELATED-cyclin-dependent kinase inhibitors (Barrada et al., 2019; Forzani et al., 2019), have further confirmed the existence of this link. It has also been shown that TOR inactivation in well-fed Arabidopsis seedlings results in the repression of expression of genes involved in the cell cycle (Caldana et al., 2013). This is consistent with the findings of Xiong et al. (2013) that carbon-starved seedlings fail to re-initiate growth after being supplied with sugar in the absence of TOR activity and that cell cycle-related genes, which are induced after 2 hours of exogenous sugar supply, require TOR activity for their expression to be induced.

In addition to the sugar status, it has been shown that the activity of the TOR complex is modulated by a broad range of signals both exogenous, like environmental stresses, and endogenous, such as plant hormones (Dobrenel et al., 2016a). While many studies have focused on the link between TOR and the sugar signaling pathway or light perception (Dobrenel et al., 2013; Pfeiffer et al., 2016; Xiong et al., 2013), recent publications have highlighted a link with other nutrients (Canellas et al., 2018; Couso et al., 2019; Dong et al., 2017). Couso et al. (2019) showed that, in *Chlamydomonas reinhardtii*, a limitation in phosphate availability results in a decreased level of LST8 protein as well as in a reduction in TOR kinase activity, and that the *psr1* (Pi starvation response1) mutant, which is defective in responses to phosphate starvation, is partially resistant to TOR inhibitors. As photoautotrophic organisms, plants use light energy to convert CO_2_ into carbohydrates in their leaves, while nutrient uptake by roots is necessary for the biosynthesis of amino acids and nucleotides coupled to energy derived from photosynthesis and respiration during the day and night, respectively. Although sulfur and glucose are assimilated in different organs, Dong et al. (2017) showed that the modulation of the TOR signaling pathway by sulfur availability is operated via the regulation of glucose metabolism. Based on this and on the meristematic expression of TOR, we hypothesize that the signaling pathway(s) resulting in the modulation of TOR activity in response to these nutritional levels may require intercellular communication.

In this paper, we focus on understanding the tight interconnection between perception of the nutritional status and modulation of TOR activity at the cellular level. We particularly focus on carbon nutrition as a source of energy and the phosphate nutrition that is indispensable for the production of nucleic acids and necessary to support growth. To do so, we took advantage of an Arabidopsis cell suspension system, which was previously successfully used to dissect the sugar signaling pathways and the establishment of photosynthesis (Dubreuil et al., 2018; Kunz et al., 2014). Employing a similar system has proven to be useful for the identification of new direct targets of the TOR complex (Van Leene et al., 2019). We believe that using a system of rapidly dividing cells mimics the meristematic TOR responses better than whole seedling or plant-based systems where meristematic activity is considerably diluted among other cell types. Moreover, the use of this system enables time-resolved experiments as well as the replicates necessary for a systems biology approach. To better understand the cellular effects of nutrient starvation and TOR inactivation, we used a systems-based approach combining metabolomics, transcriptomics and lipidomics. With this comprehensive approach, we were able to show that the TOR pathway integrates the cellular phosphate level to actively modulate the cell cycle and growth in constantly dividing cells. This is in contrast to sugar availability, which controls growth in meristematic cells through an apparently TOR-independent pathway.

## Results

### TOR inactivation, phosphate starvation or carbon starvation all result in decreased growth

Similar to other multicellular eukaryote organisms, vascular plants contain a large diversity of cell identities, ranging from dividing non-differentiated meristematic cells to highly specialized leaf mesophyll cells. This diversity complicates the study of cellular responses to nutritional and environmental signals. Cell culture systems have been used extensively in the past to study, for example, the cellular response to sucrose starvation (Nicolaï et al., 2006). More recently, a similar approach has proven to be successful in studying plant cell differentiation into tracheary elements and photosynthetic cells (Dubreuil et al., 2018; Pesquet et al., 2010) or investigating sugar signaling pathways (Kunz et al., 2014), as well as for the identification of new interactors and direct targets of the TOR kinase complex (Van Leene et al., 2019). In this study, we used a cell culture system to decipher the molecular roles of carbon and phosphorus nutrition in the regulation of growth, particularly focusing on the integration of the TOR pathway in this cellular process.

We first characterized the growth of the cell line as well as the medium composition under routine conditions of normal growth. The cells present a typical growth curve composed of a lag phase of approximately 3 days followed by an exponential growth phase finishing around the 7^th^ day of culture (which is the day on which cells are routinely subcultured) and the cells then experience a rapid decrease in their growth rate and reach maximal biomass after 10 days of culture (figure 1A). It is important to note that, although growth is not suddenly arrested after the end of the exponential phase, only a modest biomass increase is observed after days 7-8, which correlates with the complete exhaustion of sucrose from the medium (figure 1B). Biomass accumulation after the exponential growth phase may then be supported by the presence of glucose and fructose in the medium, potentially deriving from the conversion of sucrose by apoplastic invertases (supplemental figure 1), as well as by the nitrogen remaining in the medium since 1/5 of the initial pool of nitrate is still present in the medium at this stage (figure 1B).

**Fig. 1.**
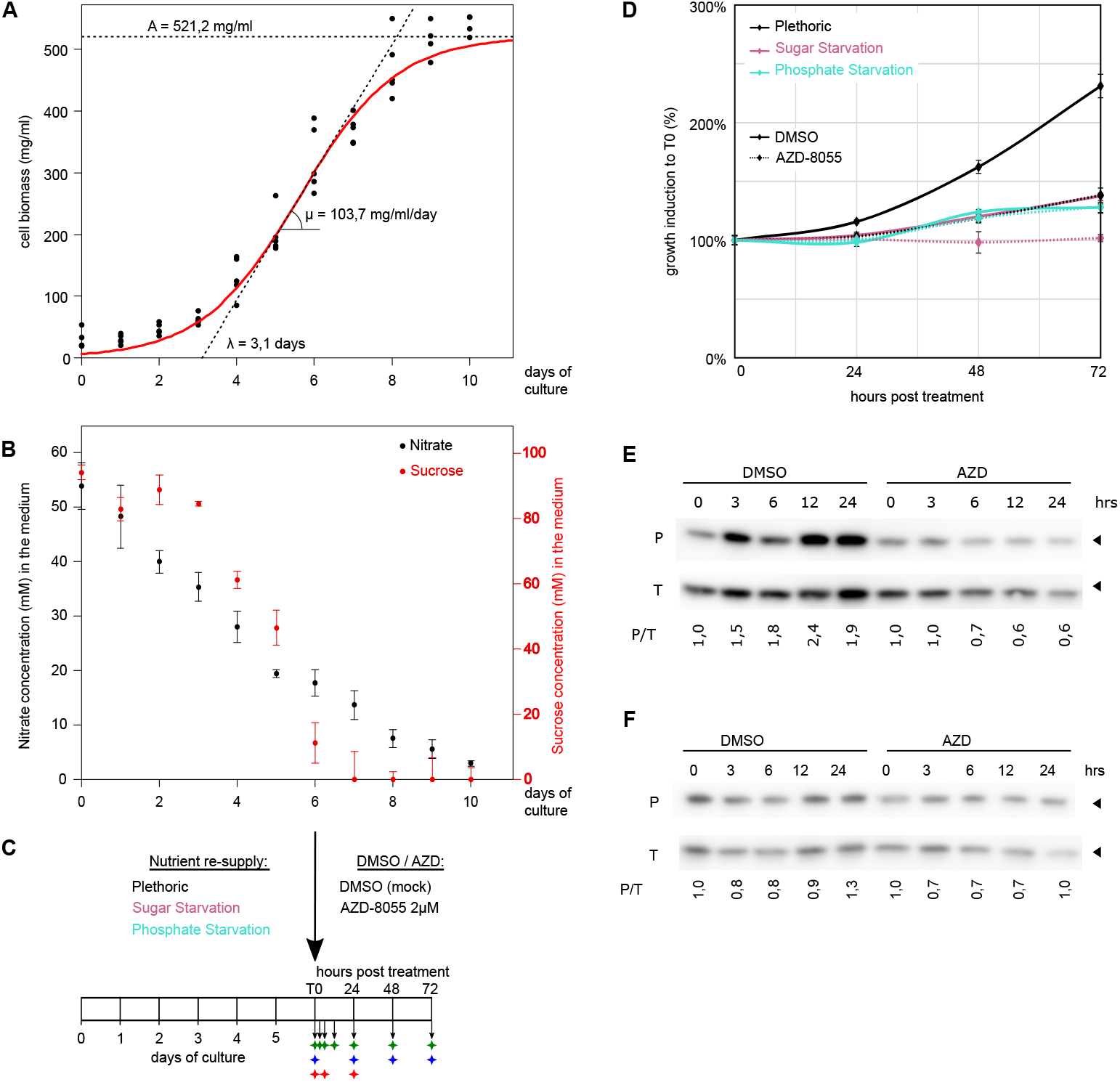
Cell culture growth parameters. **(A)** Growth curve with determination of growth parameters (A: maximal growth value, λ: period of lag and μ: maximum growth rate) calculated with the Grofit package. Dots represent experimental data and the red line represents the theoretical growth curve corresponding to the calculated parameters. **(B)** Evolution of medium composition during the cell culture growth period revealed by determination of nitrate and sucrose contents. **(C)** Experimental design: 6-day-old cells were diluted in fresh medium with or without sucrose and phosphate and concomitantly treated with 2 μM AZD-8055. Cells were harvested over a timecourse and samples obtained were analyzed by a multi-omics approach. Arrows represent harvest timepoints, green diamonds represent metabolomics and lipidomics analysis, blue diamonds represent biomass measurement, red diamonds represent transcriptomic analysis. **(D)** Evolution of biomass increase after dilution in fresh medium. **(E-F)** Evolution of RPS6 phosphorylation determined by measurement of the level of phosphorylated RPS6 (P) and of the total RPS6 level (T) by western blot analysis under plethoric conditions **(E)** or after phosphate starvation **(F)**. The phosphorylation ratio (P/T) was determined for 3 independent replicates and normalized with respect to T0. Arrowheads indicate the position of the 35 kDa marker.

For this study, we used 6-day-old cell cultures. This corresponds to the stage before the cells exit the exponential phase and initiate multi-nutrient starvation responses, although the medium is already nearly depleted of sucrose (figure 1B, supplemental figure 2, supplemental table 2). We diluted the cells at a 1:1 ratio in double strength MS medium with or without sucrose and with or without phosphorus and combined this treatment with a simultaneous application of a 2 μM concentration of the TOR inhibitor AZD-8055 (figure 1C). This created control conditions in which cells were fed with non-limiting amounts of nutrients and conditions in which cells were starved of phosphorus or sugars, as well as conditions in which TOR was chemically inactivated under plethoric or starved conditions. As expected, we observed that biomass accumulation was reduced in a sucrose- or phosphorus-free medium when compared to the plethoric conditions (figure 1D). Interestingly, this reduction in biomass accumulation was at the same level as that in the cells cultivated in the presence of AZD-8055 under plethoric nutritional conditions. However, when AZD-8055 treatment was combined with sugar starvation, growth was totally abolished, a much more severe effect than that observed after the individual sugar starvation or AZD-8055 treatments. In contrast to the combined sugar starvation and AZD-8055 treatment, the application of AZD-8055 did not have any additive effect to that of phosphorus starvation in inhibiting growth. This suggests that the regulation of growth in response to chemical inactivation of the TOR pathway and phosphorus availability may, at least partially, overlap, while in contrast, the regulation of growth by modulation of the TOR pathway and by sugar availability seem to be independent.

To assay TOR activity in the samples, we determined the level of phosphorylation of the S6 ribosomal protein (RPS6), which has been shown to be a proxy for TOR activity (Dobrenel et al., 2016b). As expected, cells subcultured with fresh medium showed increased TOR activity within the first 24 hours after the treatment (figure 1E, supplemental figure 3). Conversely, AZD-8055-treated cells showed a clear reduction in the level of phosphorylated RPS6 protein. However, increased TOR activity was not observed in phosphorus-starved cells, in which the RPS6 phosphorylation level was seemingly unaffected by AZD-8055 treatment (figure 1F, supplemental figure 3). Overall, this confirms the link between phosphorus availability and TOR activity, possibly placing perception of the phosphorus pool upstream of the modulation of TOR activity.

### TOR inactivation and phosphate starvation induce similar changes in amino acid levels

It has previously been shown that TOR inactivation in plants grown under favorable conditions induces major metabolic reprogramming, characterized mostly by a general accumulation of amino acids (Caldana et al., 2013; Dobrenel et al., 2013; Moreau et al., 2012; Ren et al., 2012). We investigated the metabolic profile of the cells after the application of AZD-8055 using a GC-MS approach (supplemental figure 4). We observed a general accumulation of amino acids in the AZD-8055-treated samples, thus confirming that the effect of TOR inactivation on amino acid accumulation in our cell culture based system is similar to that previously reported in plants.

To compare the effect of TOR inactivation with that of nutrient insufficiency, we then investigated the metabolic profile of the cells after the phosphorus and sugar limitation treatments as well as after chemical inactivation of TOR by application of AZD-8055. Hierarchical clustering of the metabolic profiles showed a clear dichotomy between the samples, with the first group containing the non-treated samples, the sugar-limited samples, and the early timepoints during AZD-8055 treatment (3 hours, independently of nutritional level) and the early timepoints of phosphorus limitation (3, 6 and 12 hours) (figure 2, supplemental table 4). In the second group, on the other hand, we found all the later AZD-8055 treated samples (independently of nutritional level) and the late samples treated with phosphorus limitation (from 24 hours of treatment onwards). This clear separation between samples is largely supported by an elevated level of amino acids after chemical inactivation of TOR and a prolonged period of phosphorus limitation. In contrast, these metabolites show decreasing levels over time in the other conditions, including the sugar-limitation treatment. Interestingly, while the amino acid contents followed the same pattern of accumulation after AZD-8055-dependent TOR inhibition and phosphate starvation, the accumulation of TCA cycle intermediates seemed to be specific to the AZD-8055 treatment (independently of nutritional level); it was not observed in response to phosphate starvation (supplemental figure 5).

**Fig. 2.**
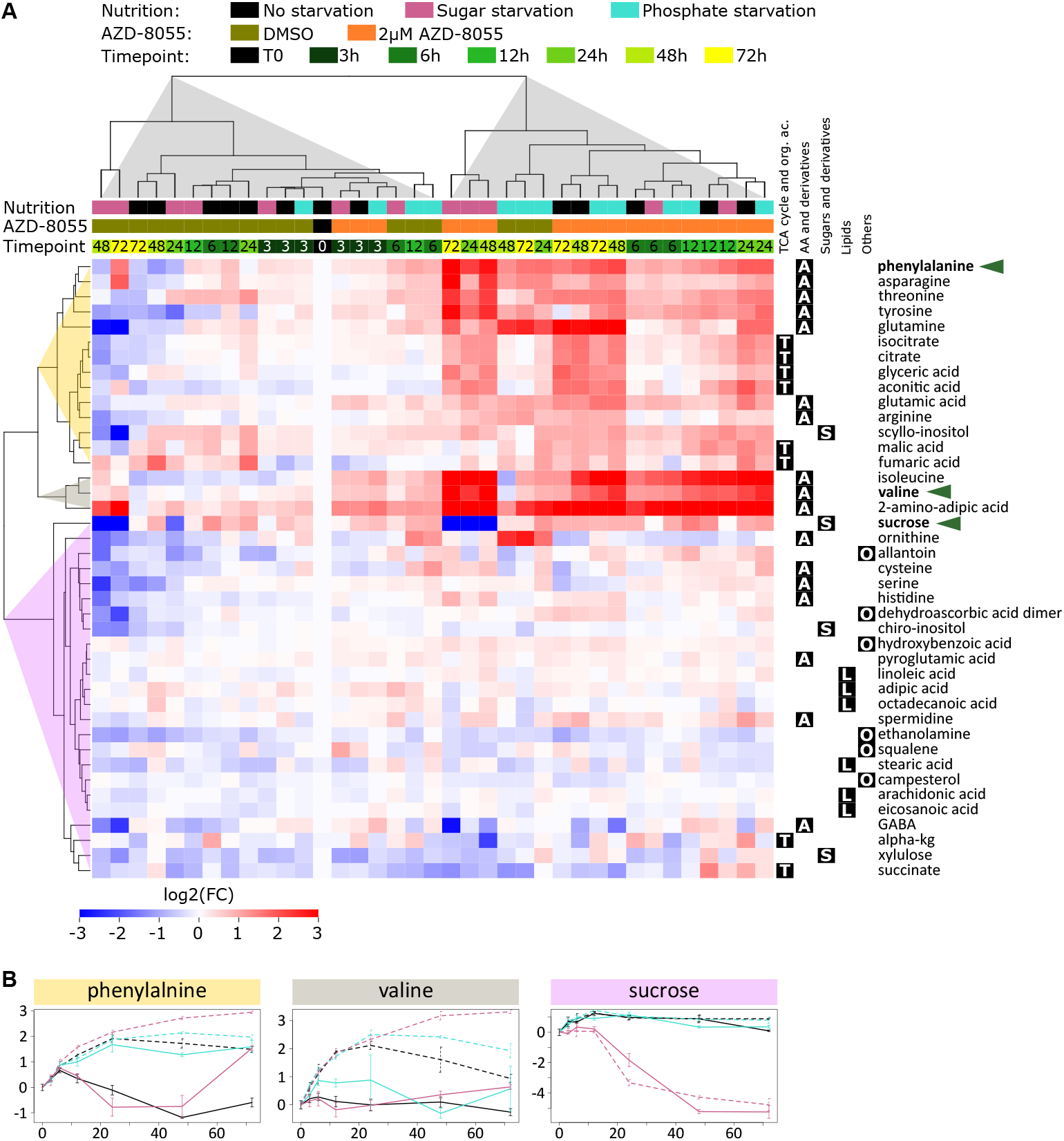
The metabolic response to TOR inactivation mimics phosphorus limitation. The metabolic profile was determined by GC-MS after phosphorus limitation, sugar limitation and/or TOR inactivation (AZD-8055). **(A)** Hierarchical clustering analysis showing the relative abundance of each metabolite over the timecourse in the different conditions after normalization to T0. The letters on the right correspond to the different metabolic categories **(B)** Evolution of relative content of 3 metabolites representative of the clusters identified in A (annotated in bold and indicated by green arrowheads). The x-axis is the time in hours and the y-axis is the abundance relative to T0 after log2 transformation. (n=3)

It is important to note that, in the sugar-limited samples, although sugar-limitation and AZD treatment were performed at the same time, the cells needed approximately 24 hours to completely exhaust their endogenous sucrose pool. This suggests that up to 24 hours, the endogenous levels of sugars were enough to maintain the pool of sucrose and that there was actual sugar starvation only from this timepoint onwards (figure 2B). We can thus divide the cellular response into two phases: the first 24 hours, corresponding to a cellular response to an exogenous limitation, and the later timepoints, corresponding to actual sugar starvation.

Based on the similarity of the response to chemical inactivation of TOR and to phosphorus limitation at the metabolomic level as well as on the absence of any additive effect between these two treatments, these data confirm the previously mentioned hypothesis of a link between phosphorus availability and TOR activity. This further suggest that TOR activity is regulated in response to phosphorus availability and that TOR regulates the level of amino acids, based on the metabolic response to TOR inactivation compared with phosphorus limitation. In contrast to phosphorus starvation, sugar starvation responses could not be linked to any increase in the amino acid level; rather there was a decrease. Indeed, prolonged carbon starvation results in depletion of most amino acids and TCA cycle intermediates, which may be possibly consumed to provide energy. Overall, this supports the hypothesis that the regulation of growth after nutrient re-supply follows two (at least partially) independent routes: one which is sugar dependent and a second which is phosphorus- and TOR-dependent.

### TOR inhibition leads to distinct shifts in the lipidome, under phosphorus or sugar starvation

Since it has been previously reported that TOR inactivation as well as phosphorus limitation induces an elevation in the level of tri-acyl glycerides (Caldana et al., 2013; Couso et al., 2019); that the *pho1* (Pi homeostasis1) mutant which is deficient in Pi transport has reduced levels of phospholipids (Rouached et al., 2011), and that phosphorus limitation leads to strong overexpression of several genes involved in lipid metabolism (Morcuende et al., 2007), we investigated the lipid content of the cell cultures during the timecourse of response to TOR inactivation in the context of nutrient limitation. We observed that the overall lipid content of the cells at the beginning of the experiment (compared to the other samples) shows elevated levels of non-phosphorus glycoglycerolipids (digalactosyldiacylglycerol DGDG and sulfoquinovosyldiacylglycerol SQDG) as well as of tri-acyl glycerides (TAG) while most of the phosphoglycerolipids (phosphatidylcholine PC, phosphatidylethanolamine PE, phosphatidylinositol PI and phosphatidylglycerol PG) were at rather low levels compared to those of the other samples (supplemental figure 6A).

Interestingly, when we followed the evolution of the lipid complement in the different conditions over time, we noticed that the nutritional level had a strong influence whereas there was only a modest effect of TOR inactivation (figure 3A, supplemental figure 6B). The samples clustered in three main groups, reflecting the nutritional status of the medium, with clear separation between the late samples of sugar starved cells in one group, the samples of phosphorus starved cells in a second group and the other treatments in a third group. The sugar-starved cells were characterized by a severe depletion in diacylglycerols and triacylglycerols (DAG and TAG), while these lipids were found at an elevated level in the phosphorus-limited cells. The phosphoglycerolipids (PC, PE, PI and PG), in contrast, showed a gradual increase over time in the absence of nutrient limitation but remained at a low level (and even tended to decrease) during phosphorus limitation. These variations are even more striking when we compare the samples timepoint by timepoint (supplemental figure 6C).

**Fig. 3.**
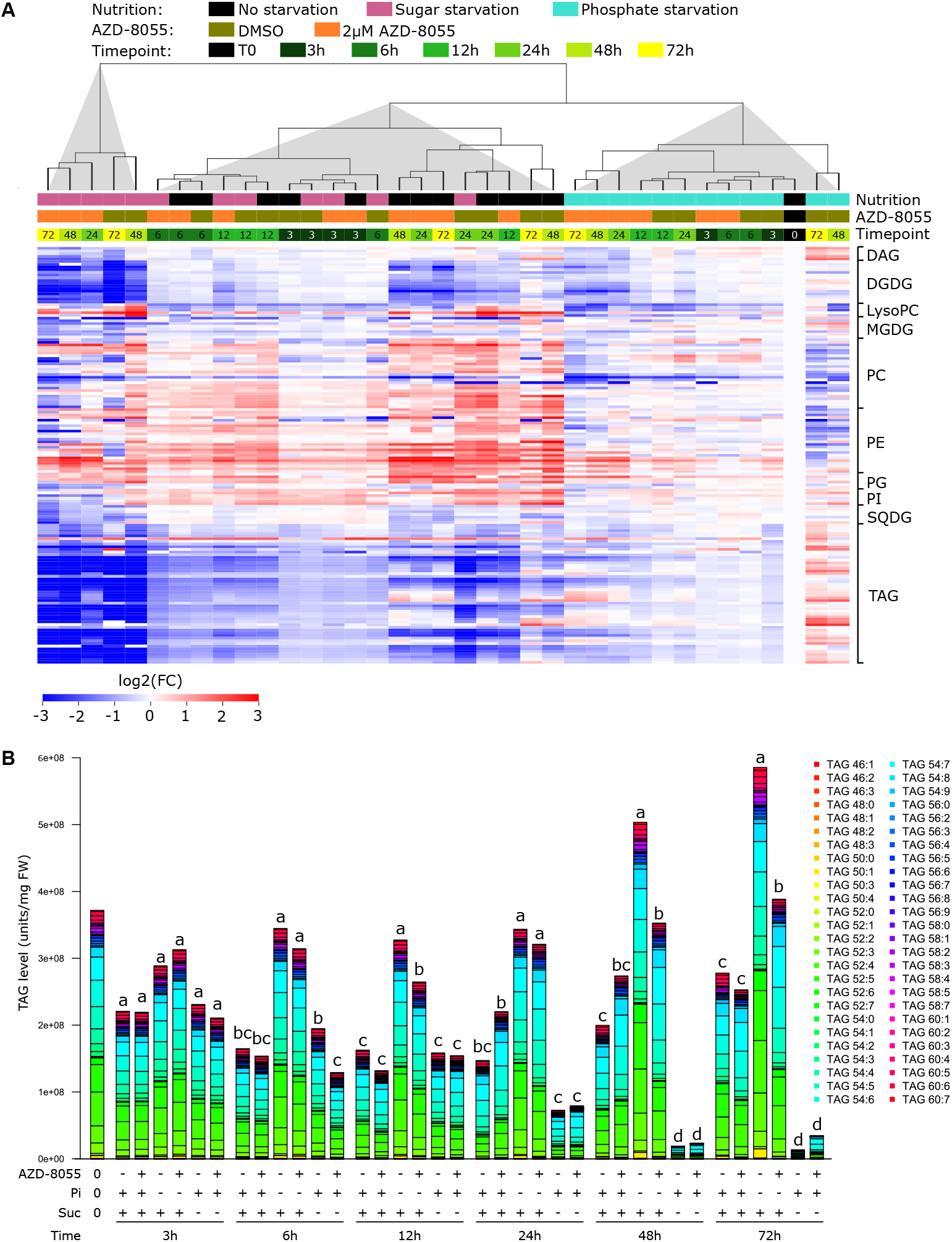
Phosphate starvation and sugar starvation have opposite effects on the accumulation of lipids, which is not affected by TOR inactivation. **(A)** Hierarchical clustering analysis showing the relative abundance of each lipid over the timecourse in the different conditions after normalization with respect to T0. The names on the right correspond to the different lipid categories **(B)** Evolution of TAG content. The letters above the histograms represent the result of an ANOVA test followed by a post-hoc Tukey analysis by timepoint (n=3)

As mentioned, the level of TAGs was strongly influenced by nutrient availability. In the absence of any nutrient limitation, the TAG level greatly decreased during the first 12-24 hours after nutrient resupply before increasing gradually (figure 3B). This late restoration of the TAG pool did not occur in the sugar-starved cells, in which the level of TAG further decreased at the late timepoints. On the other hand, in the presence of phosphorus limitation, the level of TAG remained high at the early timepoints and further increased by the late timepoints. Interestingly, contrarily to what we expected based on the literature (Caldana et al., 2013; Couso et al., 2019), TOR inactivation by application of the inhibitor AZD-8055 did not induce any increase in the TAG level; if anything it tended to repress the late increase in the TAG level after prolonged phosphorus limitation.

### Phosphate starvation acts upstream of the TOR pathway to regulate the cell cycle

As we have seen previously, there is a clear difference at both the metabolomic and the lipidomic levels between the early response to nutrient limitation and AZD-8055 treatment (3 to 12 hours) and the late response (after 24 hours). To better understand the biological processes influenced by the nutrient status and TOR at the metabolite and lipid levels as well as biomass accumulation, we performed transcriptomic analysis of the cells after 6 and 24 hours of treatment.

As expected, the re-initiation of growth stimulated by the supply of fresh nutrients was accompanied by drastic transcriptional reprogramming, with more than a third of the total transcripts being deregulated as early as 6 hours after the treatment (supplemental figure 7A, supplemental table 6). A similar number of genes was found to be deregulated after 24 hours of treatment and there was considerable overlap between the deregulated genes at the two timepoints (supplemental figure 7B-C). Among the genes that were up-regulated, we found strong overrepresentation of genes linked to cell division and the cell cycle, which must be related to the re-activation of biomass accumulation, as well as several terms linked to gene expression such as nucleotide and amino acid biosynthesis, transcription, ribosome assembly, translation. In contrast, among the genes that were down-regulated, there was a prevalence of genes linked to various stresses and programmed cell death. We also found an overrepresentation of GO terms corresponding to several metabolic pathways including lipid metabolism (supplemental figure 7D-G, supplemental material 1, supplemental table 7). Interestingly, some GO terms linked to developmental processes were found to be enriched among both induced and repressed genes at 6 and 24 hours after nutrient re-supply. Although these GO terms were present in these four datasets, they were more significantly enriched in the genes upregulated after 24 hours of growth, which contained GO terms linked to embryonic and post-embryonic development.

It is to be noted that the transcriptional reprogramming observed after the nutritional resupply also occurs in all the different conditions (supplemental figure 7A, supplemental table 6). However, this reprogramming is less extensive under phosphorus limitation, suggesting that a subset of the transcripts that are deregulated after nutrient re-supply are unaffected when phosphorus limitation is applied. When comparing the transcript profile after nutrient limitation or AZD-8055 treatment to the corresponding non-treated timepoint, up to 7000 genes were found to be differentially regulated after 6 hours of phosphorus limitation, rising to 8000 after 24 hours (supplemental figure 8A). While similar numbers were found for the AZD-8055 treated samples, there were far fewer genes that were deregulated after sugar limitation (1693 genes deregulated after 6 hours, 2286 after 24 hours). The genes deregulated after phosphorus limitation or AZD-8055 treatment showed a considerable overlap and correlation but only genes deregulated after 24 hours of sugar limitation overlapped with them (with only 3.6% after 6 hours of treatment but up to 47% after 24 hours of treatment) and showed a positive Pearson correlation, although it remained modest (figure 4A, supplemental figure 8B-C). We also compared the transcriptomic data obtained from the cell culture subjected to nutrient starvation and/or AZD-8055 treatment to the transcriptomic response of seedlings treated with AZD-8055 for 24 hours, and observed a strong Pearson correlation with the response to a 6-hour AZD-8055 treatment of cell cultures. This further validated the biological relevance of our transcriptomic data. Interestingly, we also observed a strong Pearson correlation between the response to AZD-8055 in seedlings and the response to 6 hours of phosphorus starvation in cell cultures.

**Fig. 4.**
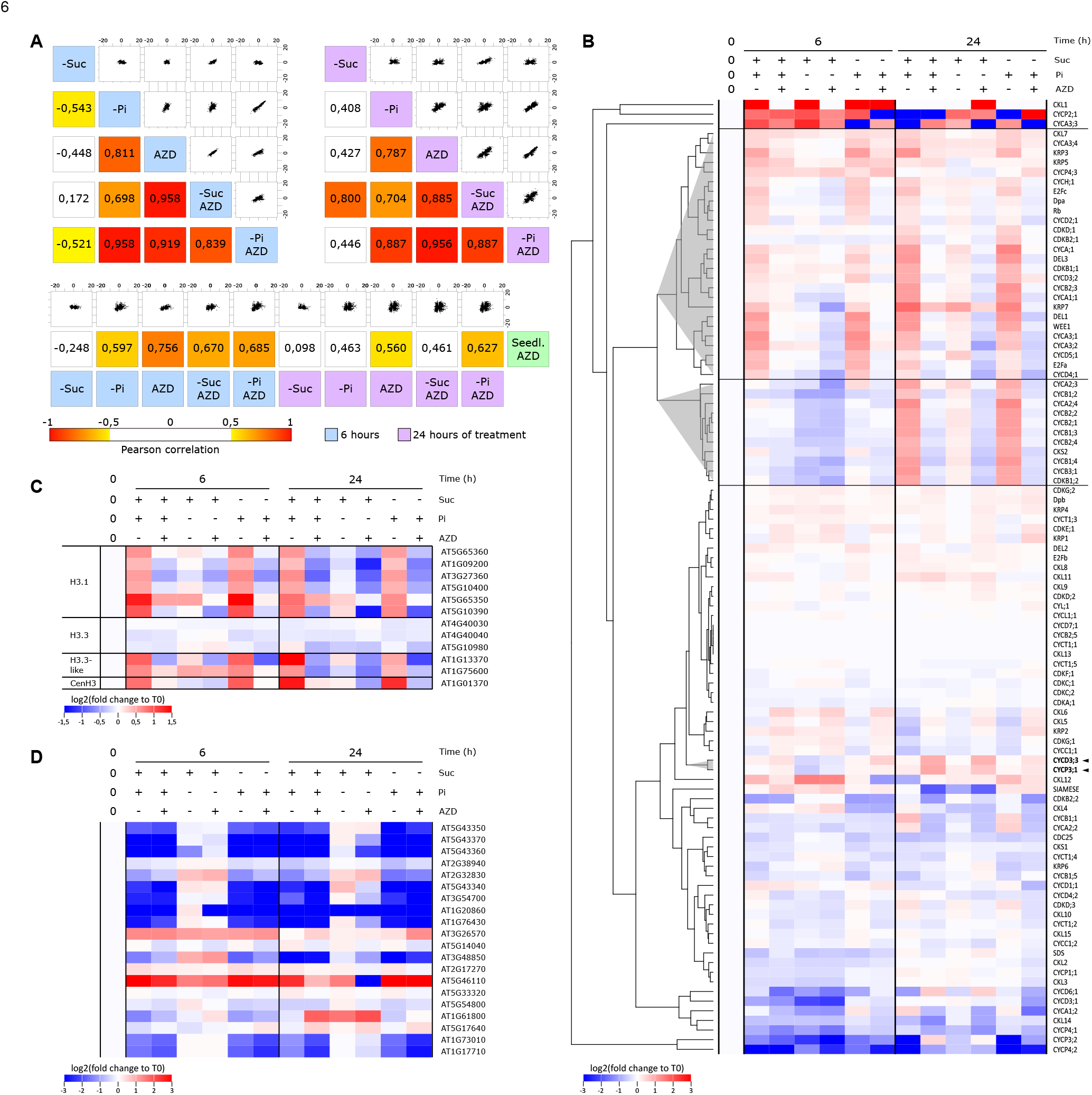
The transcriptomic response to TOR inactivation correlates with the transcriptional reprogramming after phosphorus limitation. **(A)** Correlation of transcriptional response with sugar limitation (-Suc), phosphorus limitation (-Pi) and/or TOR inactivation (AZD) after 6 hours of treatment (top left) or 24 hours of treatment (top right), and comparison with 8-day-old Arabidopsis seedlings treated with 1 μM of AZD-8055 for 24 hours (cell cultures are represented along the x-axis and seedling data along the y-axis). Only genes with a FDR < 0.01 were kept for this analysis **(B)** Expression profiles of cell cycle related genes (Menges et al., 2005) normalized to the expression profile in the T0 samples. Genes are clustered based on their responses to the different treatments **(C)** Expression profiles of phosphate transporters (Poirier and Bucher, 2002), relative to their expression values in the T0 samples **(D)** Expression profiles of H3-encoding genes, relative to their expression values in the T0 samples.

As expected based on the high number of genes overlapping between the responses to phosphorus limitation and TOR inactivation, enrichment of many gene ontology terms was shared (supplemental figure 8D, supplemental table 8). Among the induced genes, there was overrepresentation of genes linked to stresses, lipid catabolism, phosphorus metabolism and protein phosphorylation as well as carbohydrate metabolism. The repressed genes were mostly enriched for terms such as DNA conformation and organization, transcription and translation, cell cycle and embryonic development. Most of these terms were not found to be significantly enriched among the genes that were differentially expressed in response to sugar limitation. The sugar starvation induced genes were enriched for the terms response to stresses (mostly after 24 hours), cell wall organization (only after 6 hours), carbohydrate metabolism, cell growth and photosynthesis. We also observed a marked shift between the two timepoints with, for example, the response to photosynthesis being mostly enriched among the repressed genes after 6 hours but among the induced genes after 24 hours, similar to what was observed for the GO terms corresponding to carbohydrate metabolism. It is important to note here that the various terms linked to the cell cycle were not enriched among the genes deregulated after sugar limitation.

When focusing on the genes deregulated 24 hours after treatment (which is the first timepoint for which we have evidence of growth inhibition in these three treatments), we found only a few genes commonly deregulated (402 genes commonly induced and 676 genes commonly repressed) (supplemental figure 8, supplemental table 6). The up-regulated genes did not show any strongly significant GO term enrichment except for some terms linked to lipid metabolism and amino acid or organic acid metabolism (but the adjusted p-value was still quite high) (supplemental figure 8E, supplemental table 9E). In contrast, genes commonly down-regulated showed marked GO enrichment for terms linked to transcription and translation (supplemental figure 8E, supplemental table 9L) which could then be potentially linked to the absence of growth reinitiation. Interestingly, genes related to cell cycle activities were found to be commonly repressed after 24 hours of AZD-8055 treatment or 24 hours of phosphate starvation (supplemental figure 8E, supplemental table 9I). It is also worth noting that genes corresponding to the GO terms linked to phosphorus metabolism are found not only among the genes commonly induced after phosphorus starvation or AZD-8055 treatment but also among the genes specifically induced after AZD-8055 treatment (supplemental figure 8E, supplemental table 9C), suggesting that the TOR pathway plays indeed a role in the modulation of the response to phosphate starvation and that modulating it might be sufficient to mimic phosphate starvation.

In cell cultures, cell division is a key determinant of growth. We therefore focused on genes directly related to the regulation of the cell cycle (Menges et al., 2005) and observed that many of them were deregulated after some of the treatments (figure 4B). Among these genes, two groups particularly caught our attention. The first group is composed of genes that were induced in response to nutrient re-supply but repressed when it was accompanied by phosphorus limitation or TOR inactivation. It consists mainly of A-type and D-type cyclins. The second group is similar but the deregulation occurs only after 24 hours of treatment and this group contains mostly B-type cyclins. Surprisingly, the expression of these genes was unaffected by sugar limitation and starvation and their expression levels was comparable to what was observed in absence of any nutrient limitation. We also focused on the genes coding for the Histone3 protein family and observed a similar trend (figure 4C). They are induced as early as after 6 hours of nutrient resupply but not affected or repressed in response to phosphorus deficiency or TOR inactivation. Desvoyes et al. (2020) have recently shown that three genes are particularly important in idetifying the stage of the cell cycle in a given cell. Based on the expression profile of these three genes, it appears that the cell cycle is re-initiated as early as 6 hours after nutrient resupply (even in the absence of sugar in the medium) (supplemental figure 9). However, phosphate starvation and/or chemical inactivation of TOR contributes to keeping their expression at a low level.

Since it has been shown that nutritional status influences the expression of the *TOR* gene in maize (Canellas et al., 2018), we focused our attention on the expression of the genes coding for the components of the TOR complex (supplemental figure 10A). As previously reported (Moreau et al., 2010), the expression of *RAPTOR.5G* was found to be lower than that of the gene *RAPTOR.3G*. However, contrary to published work (Moreau et al., 2012), we managed to detect some expression for the gene *LST8-2* although it was thought to be a pseudogene. Its very low expression level might explain why it escaped detection in previous analysis. Interestingly, the genes coding the core TOR complex (e.g. the genes with the highest expression level, namely TOR, *LST8-1* and *RAPTOR.3G)* were found to be consistently induced after TOR inactivation by application of AZD-8055, although with a very modest fold change (supplemental figure 10B). Similar induction was also found for these three genes 6 hours after phosphorus limitation. This suggests that the repression of TOR activity in response to phosphorus limitation does not involve transcriptional repression of the expression of the genes coding for the TOR complex and that a negative feedback mechanism might exist to enhance the expression of these genes.

In order to better understand the link between the TOR pathway and the response to phosphorus starvation, we specifically investigated genes responsive to the phosphorus level (Poirier and Bucher, 2002; Thibaud et al., 2010). We observed that, as expected, the phosphate transporters were mostly transcriptionally repressed after nutrient resupply, except when it was accompanied by phosphorus starvation, conditions under which these genes are not transcriptionally deregulated, compared to T0. However, it is strikingly noticeable that regulation of their expression is seemingly unaffected by chemical inactivation of TOR (figure 4D). Interestingly, we found similar results for genes that have been previously shown to be transcriptionally deregulated in response to phosphorus limitation (supplemental figure 11), with the pattern being particularly obvious for genes that have been described as being induced by phosphorus limitation, such as genes involved in Pi recycling, signaling and sensing. The lack of response of these genes to AZD-8055 treatment highlights the existence of a phosphate-dependent but TOR-independent pathway. This further prompts us to believe that the TOR signaling pathway is recruited by the phosphate signaling pathway and is therefore downstream of it.

## Discussion

Growth in multicellular organisms is the combined result of cell expansion, including synthesis of cellular content, and cell division. It has been shown that in plants, TOR (Target Of Rapamycin), a conserved master regulator of growth in eukaryotic organisms, is expressed mostly in meristematic tissues (Menand et al., 2002). It is therefore believed that TOR controls growth through the modulation of cell division in plants (for review see Ahmad et al., 2019). This is supported by several studies showing that TOR directly phosphorylates the E2FA and E2FB transcription factors involved in the activation of the cell cycle (Li et al., 2017; Xiong et al., 2013). This link between TOR and the cell cycle was later strengthened by the characterization of the kinase YAK1 (Yet Another Kinase1) as a direct target of TOR and principal intermediate in the regulation of cell proliferation (Barrada et al., 2019; Forzani et al., 2019; Van Leene et al., 2019).

To investigate the direct effect of TOR inactivation, we therefore focused this study on a heterotrophic cell culture system where growth is mostly the result of cell division, similar to what is observed in meristematic tissues. There would be a substantial risk that, if performing the experiments in whole seedlings, the effect in cells not expressing *TOR* would overshadow the effect in meristematic tissue. As the focus of the investigation was cell proliferation, we chose to use a system of uniform and dividing cells in culture. Since TOR integrates the nutritional status to modulate growth (for review Dobrenel et al., 2016a), we compared the effect of the TOR-specific inhibitor AZD-8055 to deprivation of sugar or phosphorus, both of which have already been shown to have an effect on the modulation of the TOR pathway (Couso et al., 2019; Xiong et al., 2013). Our data suggest that, in proliferating cells, the phosphorus level is the key nutrient content integrated to recruit the TOR pathway to modulate growth.

We showed a strong overlap between the response to phosphate starvation and TOR inactivation at the transcriptomic level as well as at the metabolic level, suggesting an interconnection between these two pathways in meristematic cells. We also showed that genes involved in the primary response to phosphate starvation are unaffected by TOR inactivation while, in contrast, phosphate starvation results in repression of RPS6 phosphorylation which is a marker for TOR activity (Dobrenel et al., 2016b). This prompts us to place sensing of the phosphorus level upstream from the regulation of the TOR pathway.

In this study, we showed that sensing of phosphate starvation reduces the TOR activity level which, in turn, represses the transcription of genes necessary for re-entry into and/or progression of the cell cycle. With that in mind, we suggest that phosphate starvation (and consequently TOR inactivation) in meristematic cells actively blocks cell proliferation by regulating the expression of the cell cycle genes (figure 4B-C, supplemental figure 9). This would then repress various anabolic reactions such as protein and cell wall synthesis and DNA replication and lead to the accumulation of soluble metabolites (figure 2A), including the amino acids that are found to be accumulated in response to both phosphate starvation and AZD-8055 treatment (figure 5). Interestingly, we observed a strong overlap between our transcriptomic data and previously published transcriptomic datasets for TOR inactivation, particularly for 6 days of TOR inactivation in Arabidopsis seedlings (Caldana et al., 2013), TOR inactivation in sugar-starved seedlings (Xiong et al., 2013) and 24 hours of TOR inactivation (Dong et al., 2015) (supplemental figure 12A). This further confirms the hypothesis that the previously published transcriptomic response to TOR inactivation is based, at least for an important part of it, on a cell autonomous response, although Dong et al. (2015) already showed that there is a poor overlap between all these datasets.

**Fig. 5.**
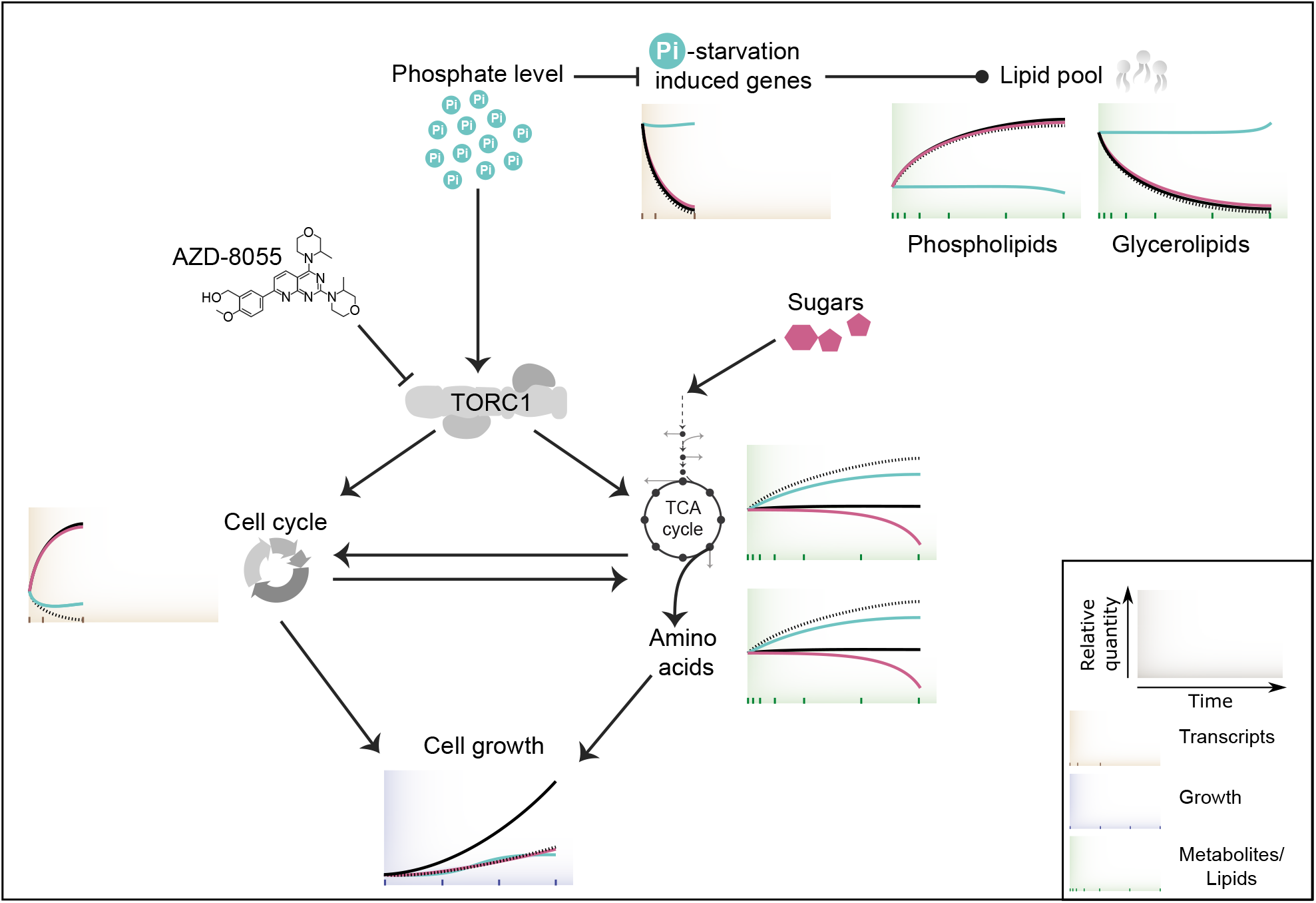
Proposed model for the phosphate TORC1 signaling pathway in proliferating Arabidopsis cells. Phosphate availability regulates TORC1 which, in turn, modulates the expression of genes involved in the cell cycle. Maintenance of cell cycle activity contributes to depletion of the TCA cycle intermediates (and further to a diminution in amino acid levels) which is fueled by the sugar pool through glycolysis. Colored boxes (brown for transcriptomic data, green for metabolomic and lipidomic data and blue for growth measurement) correspond to the representative data presented in this paper (black lines = plethoric nutrition, turquoise lines = phosphate starvation, pink lines = sugar starvation, dashed lines = AZD-8055 treatment). In the signaling cascade, arrows represent induction and activation, T-shaped arrows represent repression and the arrow with a dot at the end represents regulation.

We observed that sugar starvation has the opposite effect compared to TOR inactivation. The transcriptomic reprogramming was rather modest in comparison to the response to the other conditions. Similarly, the metabolite content was not as strongly affected as it was after AZD-8055 treatment or phosphate starvation. This weaker response may be caused by the presence of intracellular sucrose (figure 2B) up to 12 hours after the beginning of the extracellular sucrose starvation. However, it should be noted that growth limitation was observed as early as 24 hours after the treatment (figure 1D). This result is in line with another study showing that sugar starvation in cell culture results in few genes being transcriptionally deregulated although it is sufficient to arrest growth (Nicolaï et al., 2006) (supplemental figure 12B). This is surprising as Xiong et al. (2013) observed that, in sugar-deprived seedlings, exogenous application of sugars recruits the TOR pathway to re-initiate growth through a drastic transcriptomic shift. This leads us to hypothesize that sugar limitation is sensed primarily in photosynthetic tissues and requires long-distance signaling to modulate meristematic activities through the sequestration of phosphate from the meristems. This signal could possibly involve the transport of sugars from mesophyll cells to the meristems and would explain why suspension cultures which present the features of meristematic cells have lost the ability to establish a signaling response to sugar starvation.

In summary, we have shown that TOR signaling is more intertwined with the response to phosphate starvation than with the response to changed sugar levels in dividing plant cells. By using a cell culture system, where all cells are able to divide, we employ a system that resembles meristematic cells in whole plants and opens up possibilities of experiments that are otherwise impossible to perform in isolated meristems. Such an approach allowed us to measure both the transcript and the metabolite level changes over time in meristematic cells, which are where TOR expression resides, giving us access to what is most likely to be the direct response to TOR inactivation by modulating the nutrient availability in the medium or by direct pharmacological treatment. This approach may shed further light on the metabolic regulation controlled by the master regulator TOR.

## Material and Methods

### Cell culture maintenance and experiments

The cell line used is derived from root explants of *Arabidopsis thaliana* Col-0 already described (Dubreuil et al., 2018; Kunz et al., 2014). Cells were grown at 25°C in darkness with constant orbital agitation in full-strength MS salts with vitamins (Duchefa, Haarlem, The Netherlands) supplemented after autoclaving with filter-sterilized sucrose to a final concentration of 3%, and subcultured every 7 days by dilution in fresh medium at 1:10 (v/v). Cells were collected by filtration under vacuum application and extensively rinsed with ice-cold sterile distilled water. Prior to cell filtration, an aliquot of medium was harvested for analysis (see below). Cell density was calculated by measuring the fresh weight of the filtered cells and cell samples were immediately snap-frozen in liquid nitrogen.

For the nutrient resupply and starvation experiments, 6-day old cells were diluted at 1:1 in 2X minimal Murashige and Skoog medium (containing only vitamins and micronutrients) and macronutrients were provided as indicated in the supplementary table 1. In short, the medium used contained, or did not, a final concentration (after dilution by the cells) of 3% sucrose and contained, or did not, 1.25 mM of KH2PO4. AZD-8055 dissolved in DMSO was added to some cultures immediately after the addition of macronutrients, to a final concentration of 2 μM, while an equivalent volume of DMSO was used as a mock treatment.

### Determination of medium composition

An aliquot of medium harvested before cell filtration was cleared by centrifugation and immediately snap frozen in liquid nitrogen. The total nitrate content was determined by spectrophotometry according to Hood-Nowotny et al. (2010). The soluble sugar content (glucose, fructose, sucrose) was determined by enzymatic assay according to Stitt et al. (1989).

### Plant material and growth conditions

Arabidopsis thaliana wild type Col-0 (reference N60000) seeds were used. Plants were grown on half-strength Murashige and Skoog medium with vitamins and MES buffer at pH 5.9 and 10 g/L plant agar (Duchefa, Haarlem, NL) for 8 days under long day (16h light/8h dark) fluorescent light (100 μE.m^-2^, s^-1^) conditions. Seeds were chlorine gas sterilized for four hours and subsequently pipetted onto solid growth medium using 0.1% agar solution before stratification in the dark at 4°C for 2 days. Plants were grown for 7 days before 24 hour treatment with 1 μM AZD-8055 or mock treatment. The 8 day old seedlings were harvested and snap frozen in liquid nitrogen.

### Metabolomic profiling and lipid quantification

#### Metabolomic profiling

Samples from three independent biological replicates, with two technical replicates each, were subjected to metabolomic profiling according to Kusano et al. (2011). In short, metabolites were extracted from 25 mg of frozen cell powder in a solvent composed of chloroform:methanol:water (3:1:1) containing stable isotopes of reference metabolites (see Kusano et al. (2011) for the list). Extracted metabolites were then derivatized with methoxyamine hydrochloride in pyridine and MSTFA and finally diluted in heptane. Peaks were acquired with a GC-TOF-MS according to Law et al. (2018) and retention indices were calculated based on separation of an n-alkane (C_8_-C_40_) series. Metabolites were automatically detected from the chromatograms based on annotation of an in-house UPSC library and *a posteriori* manually curated against the UPSC library and the Golm library (Schauer et al., 2005). The matrix of peak areas obtained for the confirmed metabolites was normalized for instrument sensitivity (by calibrating with the internal standards) and with respect to the fresh weight. Statistical analysis were performed with the SIMCA 13.0.3 and RStudio v. 1.2.5019 (using R v. 3.6.2) software packages.

#### Lipid extraction and UPLC/MS analysis

Lipid extraction and analysis were performed for three independent biological replicates as previously described (Salem et al., 2016). Briefly, lipids were extracted from 20 mg of homogenized tissue by suspending the material in 1 ml of pre-cooled (−20°C) MTBE extraction solution (methanol:methyl tert-butyl-ether [1:3; v/v]) spiked with 0.5 μg.ml^-1^ of 1,2-diheptadecanoyl-sn-glycero-3-phosphocholine. The samples were incubated for 30 min on an orbital mixer at 4°C followed by sonication for 10 min in an ice-cooled sonication bath. After addition of 500 μl of methanol:water (1:3, v/v) to induce phase separation, the samples were vortexed and centrifuged for 5 min at 20,000×g at 4°C. An aliquot of 500 μl was collected from the upper phase containing the lipids and dried in a vacuum concentrator. The pellet was resuspended in 250 μl acetonitrile:2-propanol (7:3, vol/vol) of which 2 μl was subjected to UPLC/MS analysis (Salem et al., 2016).

#### Lipid annotation and statistical analysis

The UPLC/MS data were processed using ToxID (Version 2.1.2; Thermo) with mass error 15 ppm, retention time (RT) window 0.05 minutes. To remove noise and contaminants, data for every lipid species with an average peak height lower than the average peak height of the method blanks or with 50% of the values below 1,000 arbitrary counts were removed from the dataset. The remaining peaks were then assigned to annotated lipid species using an in-house-generated lipid database for Arabidopsis (Hummel et al., 2011). The data were normalized to the internal standard and with respect to sample fresh weight. For some lipid species, more than one peak was detected with the same m/z and identical adducts but different retention times. In these cases, we added the letter A, B, C, or D to the compound name, depending on their elution order.

### Transcriptomics analysis

#### Treatment of cell samples

Total RNA was isolated using a QIAGEN RNeasy Plant Mini Kit according to the manufacturer’s instructions. RNA was quantified with a Nanodrop ND-100 spectrophotometer, and RNA quality was assessed using an Agilent 2100 bioanalyzer (Agilent Technologies). Libraries were prepared with an Illumina TruSeq Stranded mRNA kit with poly-A selection. For the sequencing, clustering was done by ‘cBot’ and samples were sequenced on NovaSeq6000 (NovaSeq Control Software 1.6.0/RTA v3.4.4) with a 2×151 setup using ‘NovaSeqXp’ workflow in an ‘S1’ mode flowcell. Bcl to FastQ conversion was performed using bcl2fastq v2.19.1.403 from the CASAVA software suite. The quality scale used was Sanger / phred33 / Illumina 1.8+. Data pre-processing was performed following the guidelines described in Epigenesys^1^. Briefly, the quality of the raw sequence data was assessed using FastQC^2^, v0.11.4. Residual ribosomal RNA (rRNA) contamination was assessed and filtered using SortMeRNA (v2.1; Kopylova et al., 2012) with the settings --log --paired_in --fastx—sam --num_alignments 1 and using the rRNA sequences provided with SortMeRNA (rfam-5s-database-id98.fasta, rfam-5.8s-database-id98.fasta, silva-arc-16s-database-id95.fasta, silva-bac-16s-database-id85.fasta, silva-euk-18s-database-id95.fasta, silva-arc-23s-database-id98.fasta, silva-bac-23s-database-id98.fasta and silva-euk-28s-database-id98.fasta). Data were then filtered to remove adapters and trimmed for quality using Trimmomatic (v0.39; Bolger et al., 2014, settings TruSeq3-PE-2.fa:2:30:10 SLIDINGWINDOW:5:20 MINLEN:50). After the two filtering steps, FastQC was run again to ensure that no technical artefacts had been introduced. Read counts were obtained using salmon (v0.14.1, Patro et al., 2017) with non-default parameters −gcBias −seqBias and using the ARAPORT11 cDNA sequences as reference (retrieved from the TAIR resource; Berardini et al., 2015; Cheng et al., 2017). The salmon abundance values were imported into R (R Core Team, 2019) using the Bioconductor (v3.10; Gentleman et al., 2004) tximport package (v.1.12.3; Soneson et al., 2015). For data quality assessment (QA) and visualization, the read counts were normalized using a variance stabilizing transformation as implemented in DESeq2. The biological relevance of the data - e.g. similarity of biological replicates - was assessed by Principal Component Analysis (PCA) and other visualization methods (e.g. heatmaps), using custom R scripts, available at https://github.com/nicolasDelhomme/arabidopsis-nutrition-tor. Statistical analysis of gene and transcript differential expression (DE) between conditions was performed in R using the Bioconductor DESeq2 package (v1.26.0; Love et al., 2014), with the model ~ Conditions using the T0 data as reference to account for the timepoint, the nutrition supplied and TOR inactivation or with the model ~ Conditions using the non-treated timepoint data as reference to account for the nutrition supplied and TOR inactivation at a given timepoint. FDR adjusted p-values were used to assess significance; a common threshold of 1% was used throughout together with a cutoff for logarithmic expression fold change set at 0.5 to identify deregulated genes, as suggested by Schurch et al. (2016). All expression results were generated in R, using custom scripts.

Gene ontology enrichment was performed with the BINGO plugin of Cytoscape (v. 3.7.2) and further analyzed and displayed in Gephi (v. 0.9.2) or in R with the ggplot2 package.

#### Treatment of the seedling samples

A TRIzol® Plus RNA Purification Kit from Thermo Fisher Scientific was used to isolate RNA from frozen seedling samples according to the protocol provided. The RNA samples were DNase I treated to remove remaining genomic DNA and cleaned up with the PureLink™ spin columnbased RNA isolation technology (Thermo Fischer).

RNA samples were sequenced on an Illumina HiSeq 2500 instrument using a standard protocol by Macrogen (Korea). RNA libraries were made using Illumina TruSeq technology and standard protocols. Raw sequencing reads were aligned to the Arabidopsis genome (TAIR10) using TopHat v2.0.13 (Trapnell et al., 2009) with the parameter settings: ‘bowtie1’, ‘no-novel-juncs’, ‘p 6’, ‘G’, ‘minintron-length 40’, ‘max-intron-length 2000’. On average 97.0% (94.0 – 98.1%) of the raw reads could be aligned to the genome per biological replicate. This represents an average of 51.1 (40.7 – 73.3) million mapped reads. Aligned reads were summarized over annotated gene models using HTSeq-count v0.6.1 (Anders, 2010)^3^ with settings: ‘-stranded no’, ‘-i gene_id’. Sample counts were depth-adjusted by means of the statistical computing environment R (R Core Team, 2013) using the median-count-ratio method available in the DESeq R package version 3.2.5 and the DESeq2 package version 1.4.5 (Anders and Huber, 2010).

For each comparison of three test replicate samples and three control replicate samples, a DeSeq dataset of raw counts for all 33540 genes in the 6 samples of interest was generated. Log2 fold changes and Benjamini Hochberg FDR adjusted p-values were generated using the DeSeq function from the DESeq2 package, with cooksCutoff set to false.

Lists of genes with significantly changed expression were generated using an absolute log2 fold change set at above 0.5 and a Benjamini Hochberg FDR adjusted p-value of below 0.01 as cutoffs.

### Western blots

For quantification of the phosphorylation status of the RPS6, total protein extracts were subjected to western blot analysis as described in Dobrenel et al. (2016b) using the primary antiphospho-RPS6 antibody generated for this article as well as a primary antibody directed against mammalian RPS6 (Cell Signaling Technology #2317S). For the western blots, protein contents were normalized using the primary anti-tubulin antibody (Agrisera AS10 680).

## Additional information

The transcriptomic data are available from the European Nucleotide Archive (ENA^4^) under the accession number PRJEB42585. Workflows and scripts to preprocess and analyze the data have been made available in a git repository with the DOI 10.5281/zenodo.4608028^5^.

## Acknowledgements

The authors would like to thank the Swedish Metabolomics Center (SMC), and especially Jonas Gullberg, Inga-Britt Carlsson, Krister Lundgren and Hans Stenlund, for their excellent technical assistance. The authors are also thankful to Änne Michaelis (MPIMP) for the excellent technical assistance for lipid profiling analysis. We are also grateful to the Umeå Plant Science Centre Bioinformatics facility (UPSCb) for their help regarding the transcriptomic analysis. The authors acknowledge support from the National Genomics Infrastructure in Stockholm funded by Science for Life Laboratory, the Knut and Alice Wallenberg Foundation and the Swedish Research Council, and SNIC/Uppsala Multidisciplinary Center for Advanced Computational Science for assistance with massively parallel sequencing and access to the UPPMAX computational infrastructure. We gratefully acknowledge Daria Chrobok (DC SciArt) for her help in preparing the figures. This work was supported by grants from the Knut and Alice Wallenberg Foundation and the Swedish Governmental Agency for Innovation Systems. We thank Bio4Energy, a Strategic Research Environment appointed by the Swedish government, as well as the Kempe foundation, for supporting this work.

## List of supplemental material

The supplemental tables are available upon request at

**Supplemental Figure 1.**
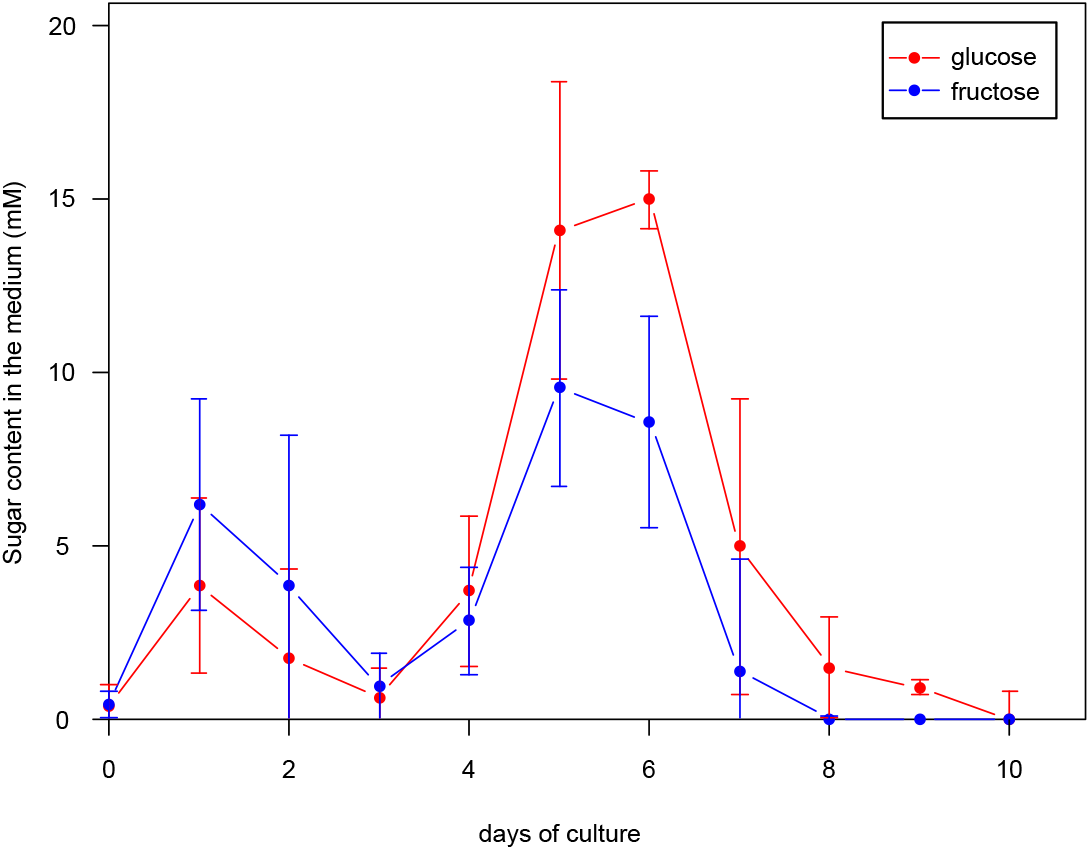
Evolution of glucose and fructose content in the cell culture medium during the growth period.

**Supplemental Figure 2.**
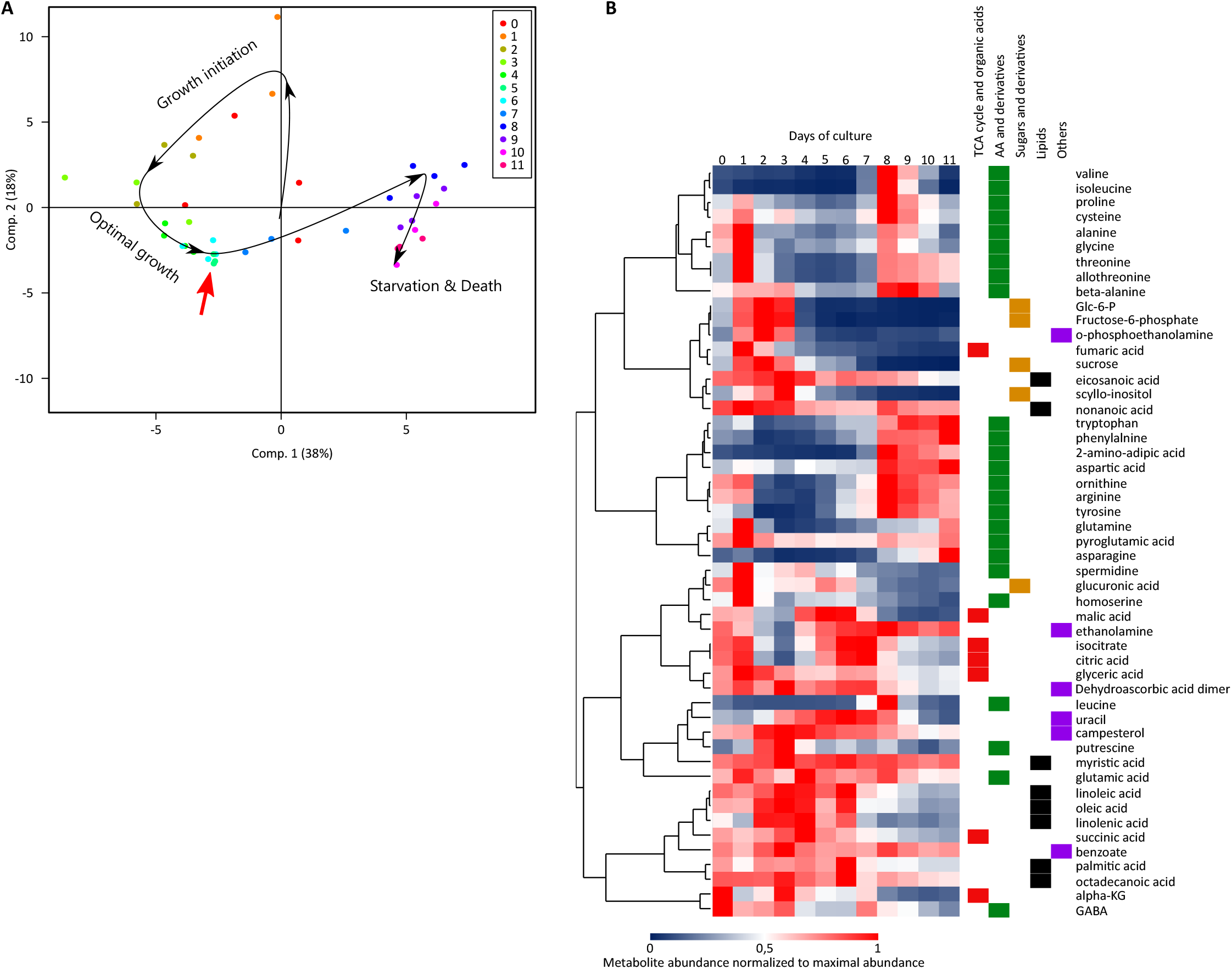
Analysis of metabolite profiles identifies three major phases occurring during the growth of a culture. (A) Principal Component Analysis showing evolution of the metabolomic profile of the cells highlighting gradual changes over the timecourse (black arrow) as well as the switch between the optimal phase of exponential growth and the beginning of starvation (red arrow). (B) Cluster analysis of metabolite content during the timecourse of growth. The relative metabolite content was measured in cells during progression of the culture starting from the subculture point (day 0), averaged per day and normalized with respect to the maximal abundance. Metabolites are separated into categories represented by the color coding on the right. (n = 4)

**Supplemental Figure 3.**
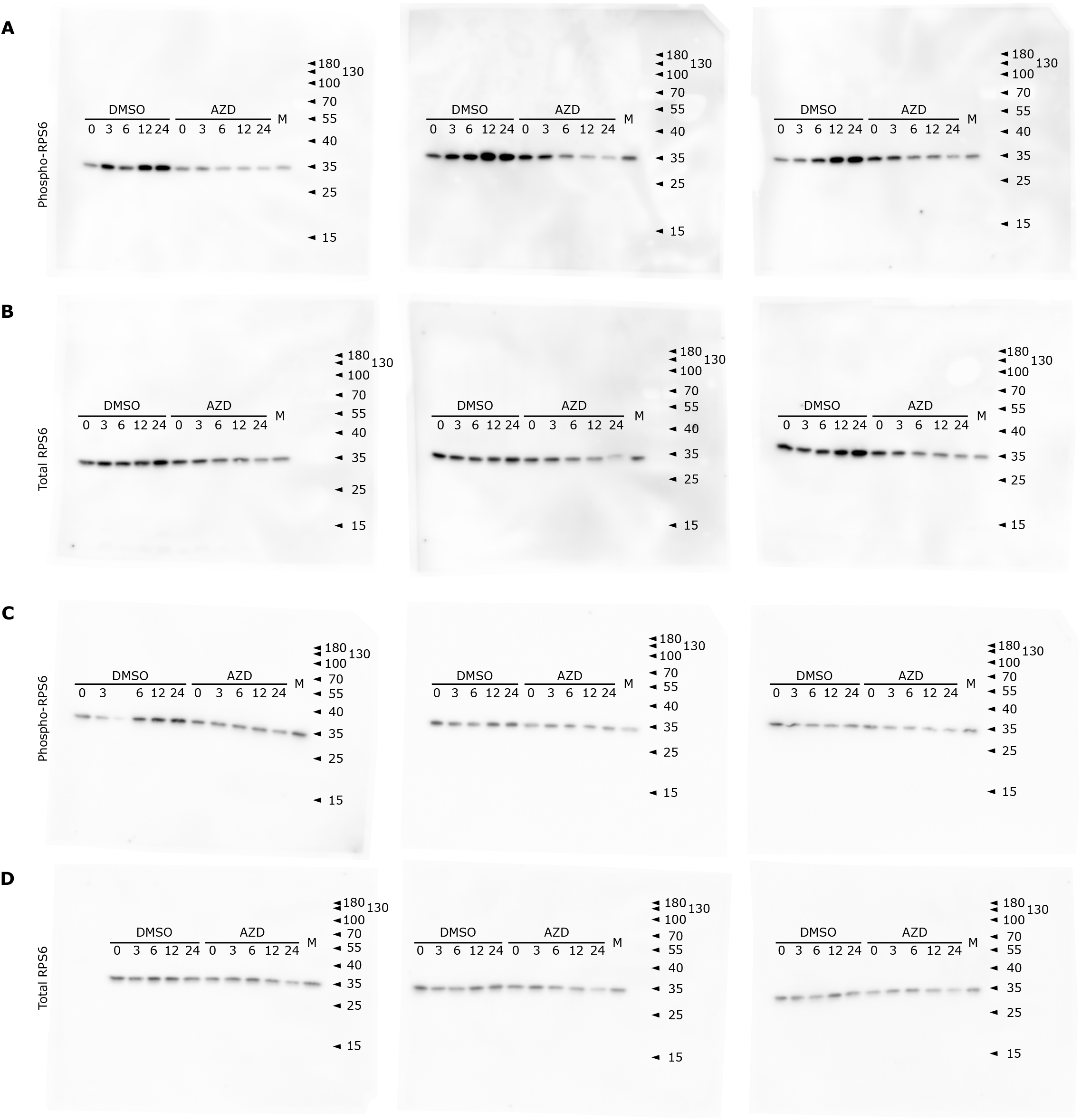
Evolution of the RPS6 phosphorylation by measurement of the phosphorylated RPS6 level (A and C) and of the total RPS6 level (B and D) by western blot under plethoric conditions (A-B) or after a phosphate starvation (C-D). Each gel picture corresponds to a different biological replicate. Numbers above the bands correspond to hours after treatment. Numbers on the right hand correspond to size markers (kDa). M is a mixed sample used for quantification standardization.

**Supplemental Figure 4.**
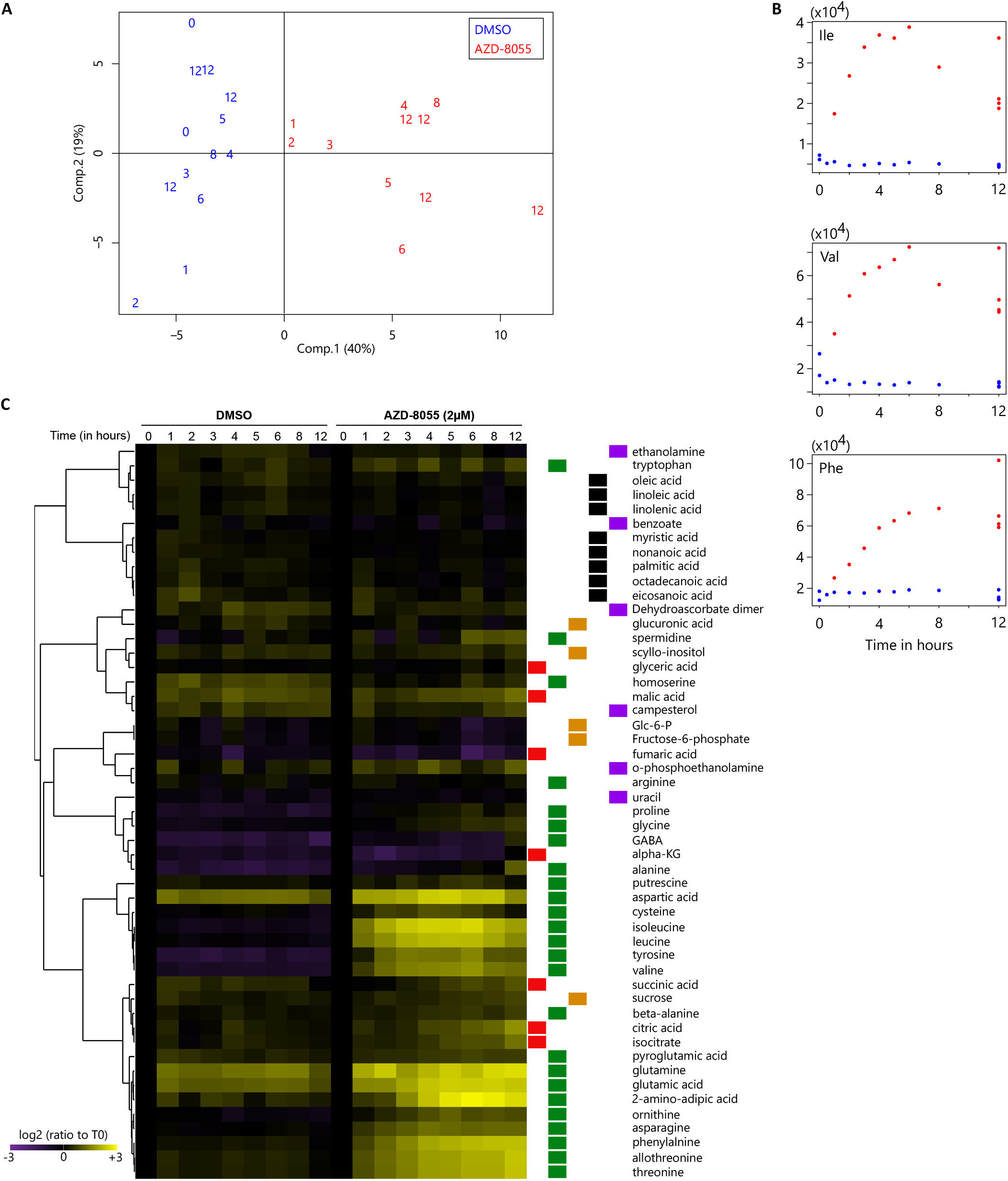
Metabolomic reprogramming after TOR inactivation in 4-day-old exponentially growing cells. (A) Principal Component Analysis plot for the metabolite profile at different time points of 2 μM AZD-8055 treatment (in red) or mock treatment (in blue) (numbers correspond to the time after treatment in hours) (B) Evolution of the concentrations of three selected amino-acids (y-axis in arbitrary units) (C) Hierarchical clustering analysis of metabolite changes during the kinetics of TOR inactivation. Relative metabolite contents are normalized to T0. The color coding on the right represents the metabolic categories (red for TCA cycle and organic acids, green for amino acids and derivatives, orange for sugars and derivatives, black for lipids and purple for others).

**Supplemental Figure 5.**
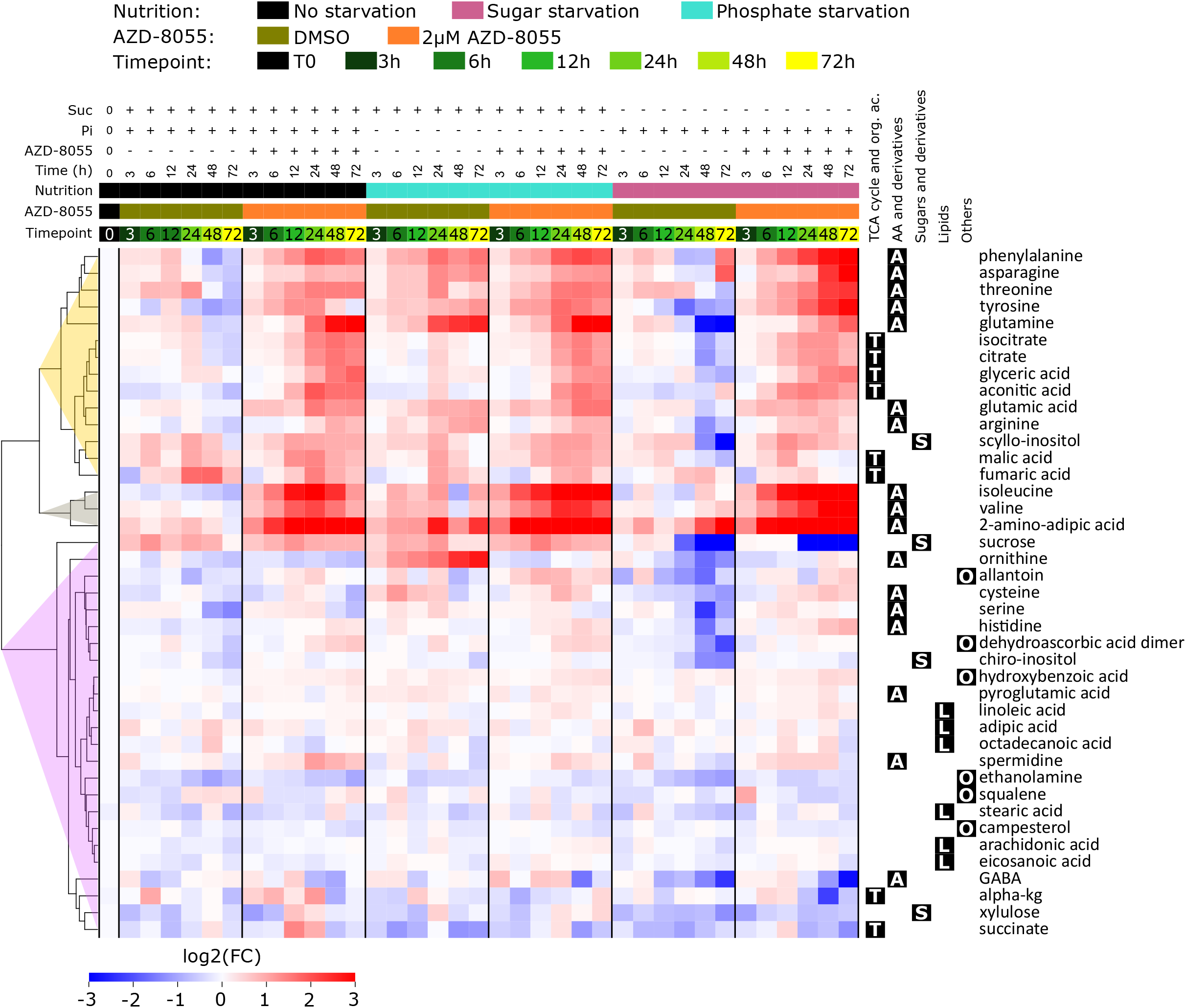
Hierarchical clustering analysis showing the relative abundance of each metabolite over the timecourse under different conditions after normalization to T0; similar to figure 2 but with the samples separated by treatment levels.

**Supplemental Figure 6.**
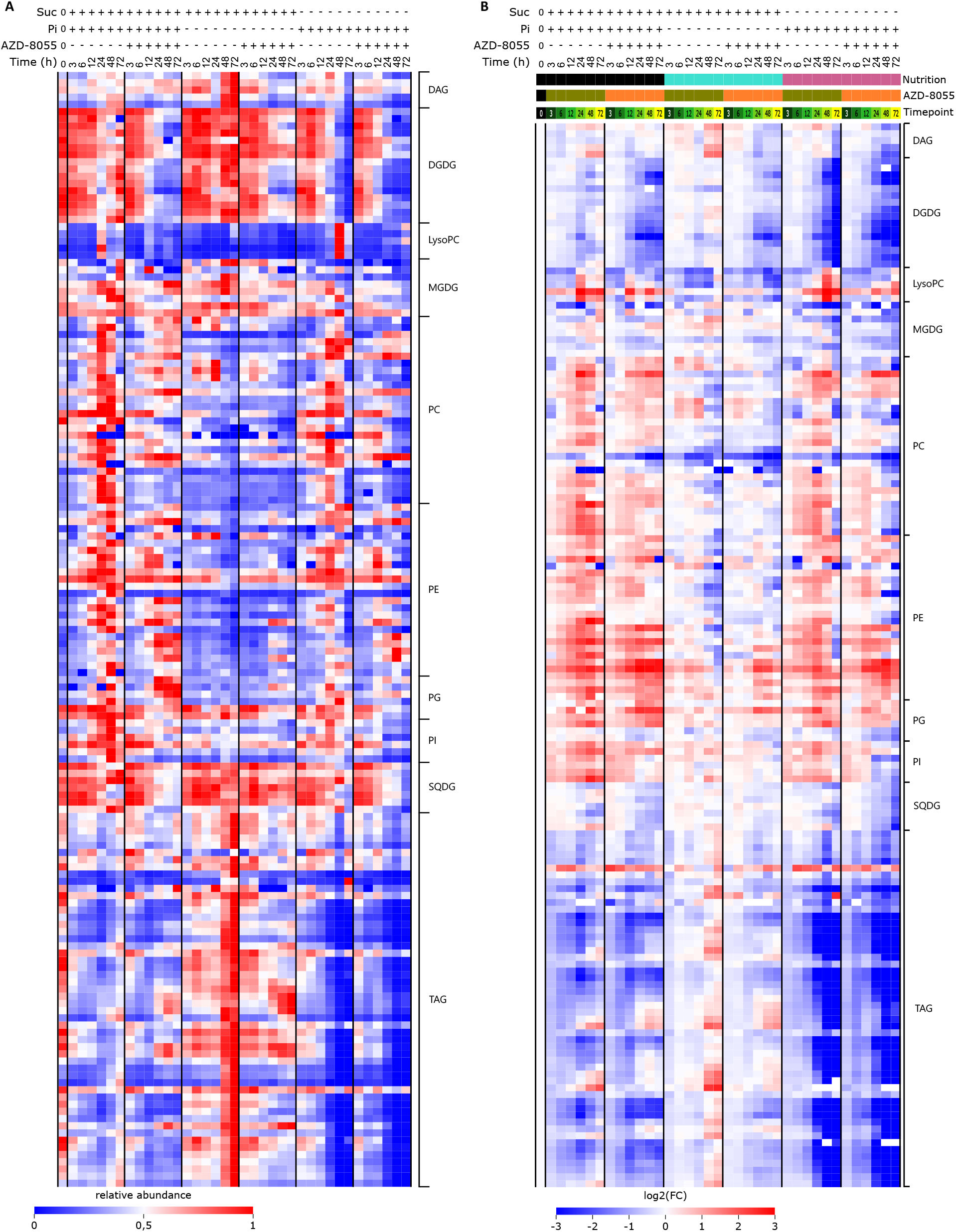
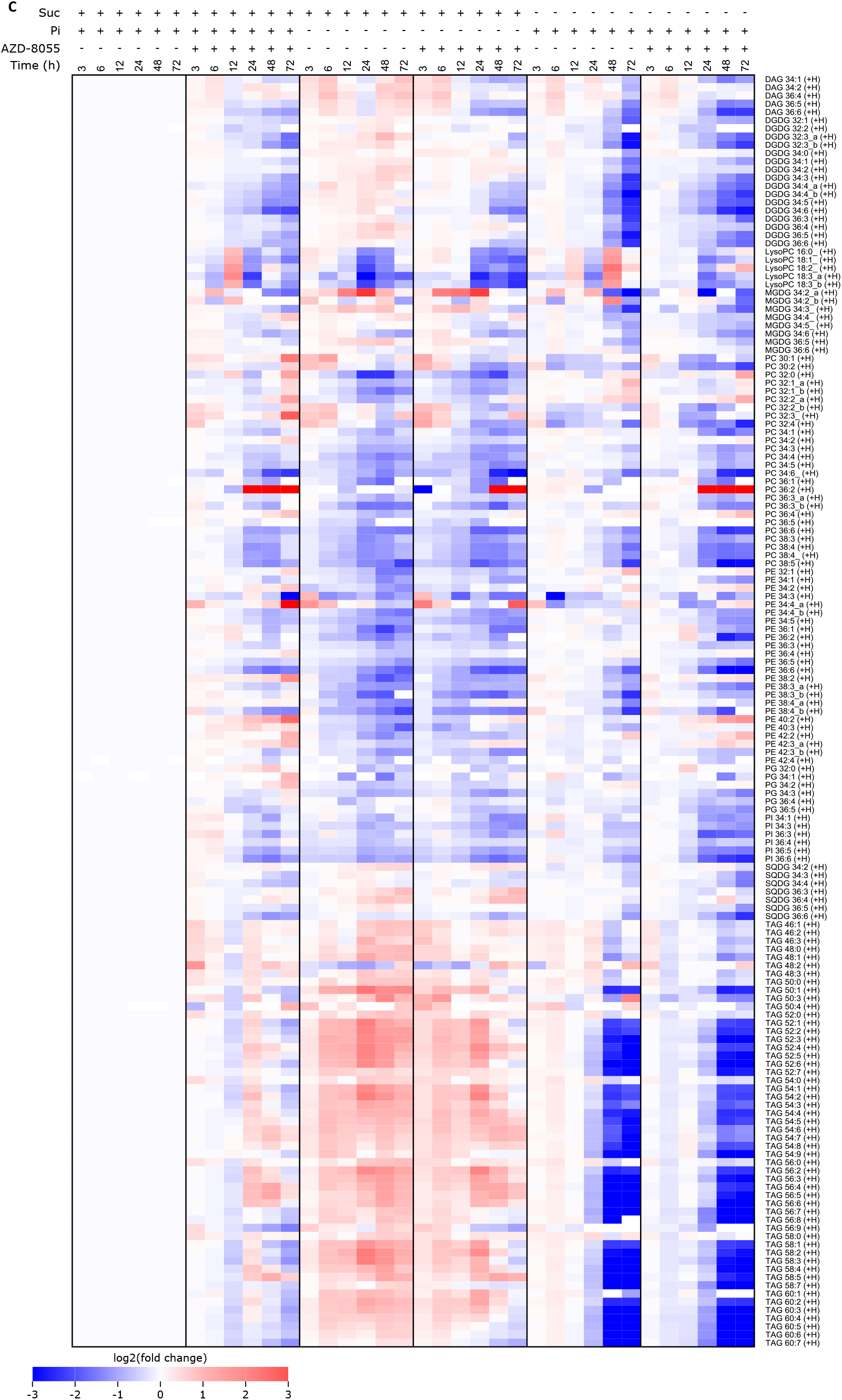
Summary of the lipidomic profile of the cell culture samples during TOR inactivation and nutrient limitation. (A) For each lipid, the values are normalized with respect to the maximal value. (B) For each lipid, the values are normalized to T0 (for the color coding of the conditions, see figure 3A). (C) For each timepoint, the values are normalized to the corresponding untreated sample (values depicted are log2 transformed fold changes).

**Supplemental Figure 7.**
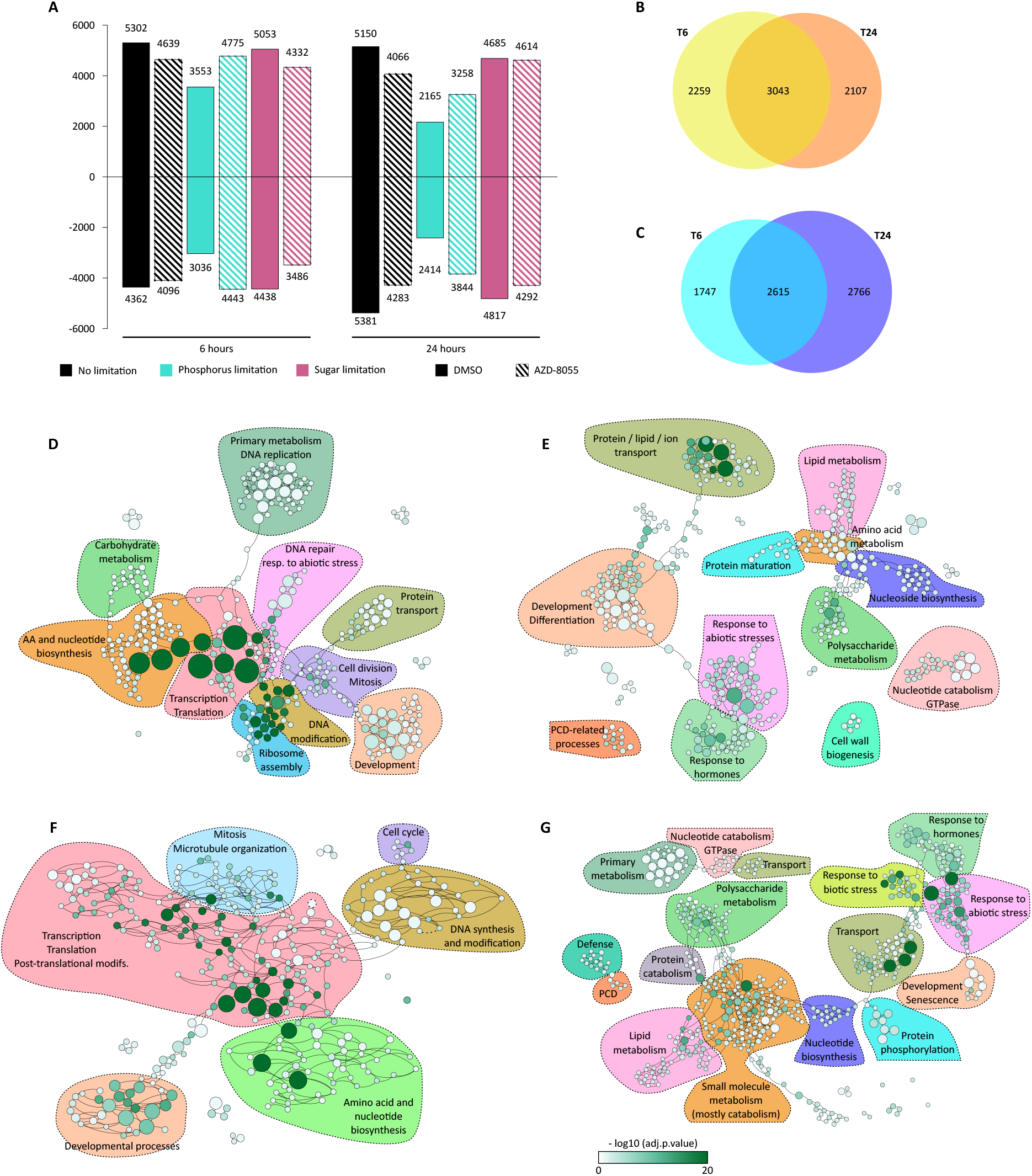
Growth re-initiation after nutrient re-supply is accompanied by extensive transcriptomic reprogramming. (A) Number of genes transcriptionally deregulated compared to the beginning of the experiment (B-C) Comparison of genes upregulated (B) or downregulated (C) after 6 or 24 hours of nutrient re-supply. (D-G) Gene Ontology term enrichment of the biological processes affected after 6 hours (DE) or 24 hours (F-G) of nutrient re-supply. (D and F) Induced genes, (E and G) repressed genes. Colors of the GO bubbles correspond to the log transformed adjusted p-values and their sizes correspond to the number of occurrences in the dataset. Only terms with an adjusted p-value > 0,05 were kept for the analysis. To improve visualization, GO terms with more than 2000 occurrences in the genome were removed.

**Supplemental Figure 8.**
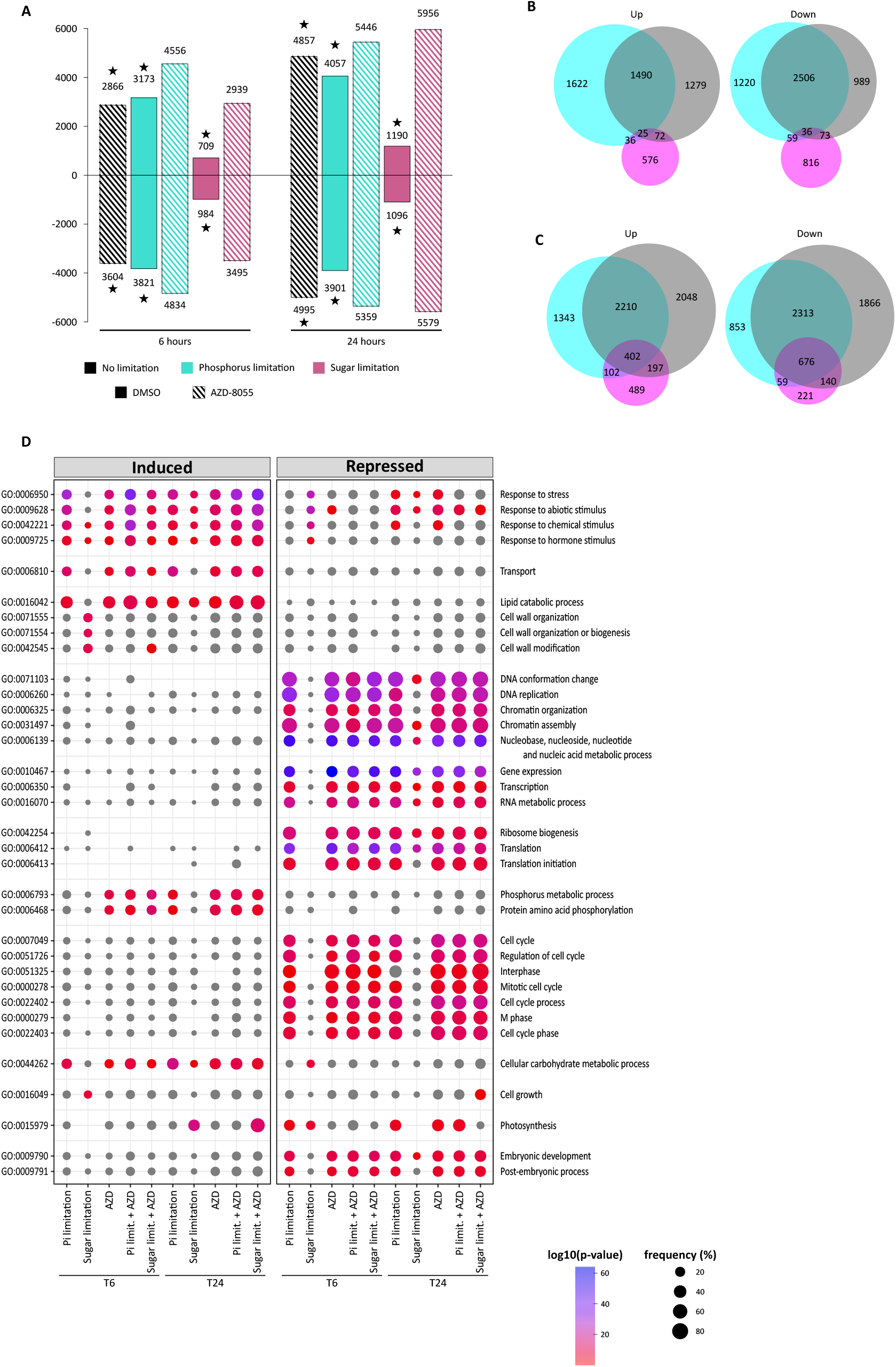
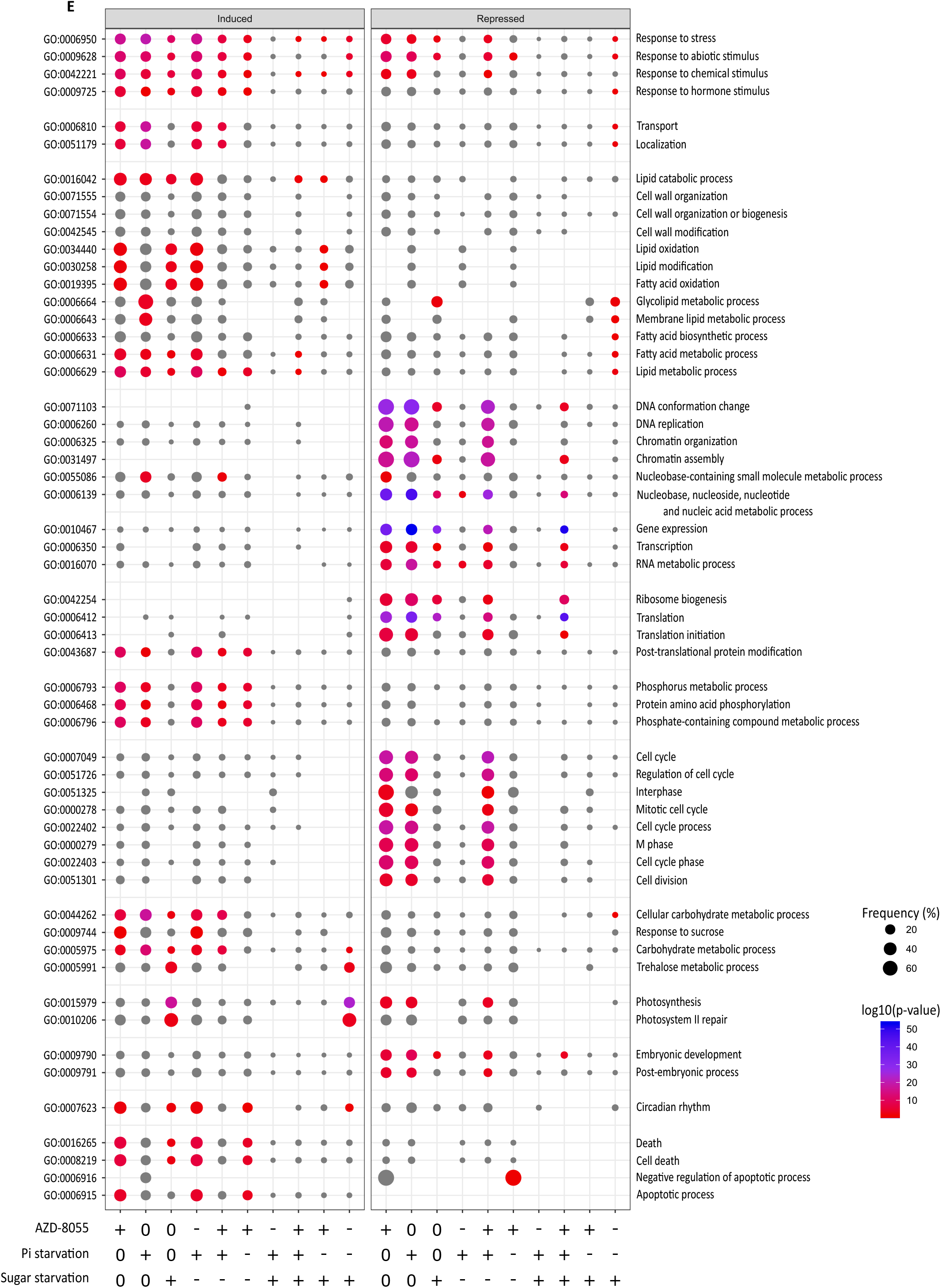
Nutritional limitation or TOR inactivation result in considerable transcriptomic reprogramming. (A) Number of genes transcriptionally deregulated compared to the corresponding non-treated timepoint. Bars above zero represent upregulated genes and bars below zero represent downregulated genes. Numbers associated with each bar correspond to the actual numbers of deregulated genes. Stars correspond to the gene sets used for panels B and C. (B-C) Venn diagrams showing the overlap of deregulated genes between different conditions after 6 hours of treatment (B) and 24 hours of treatment (C) when compared to the corresponding non-treated timepoints. Turquoise = phosphorus limitation, pink = sugar limitation, black = AZD-8055 treatment. (D) Bubble plot of the Gene Ontology terms enriched in the deregulated genes represented in panel A. Gray bubbles represent GOs with an adjusted p-value > 0,05, other colors are on a scale (shown below) indicating the log transformed adjusted p-value. The sizes of the bubbles correspond to the number of occurrences in the dataset normalized to the number of occurrences in the genome. (E) As in D for the different overlaps among the Venn diagrams presented in C (+ when the condition is included, - when the condition is excluded, 0 when the condition is ignored).

**Supplemental Figure 9.**
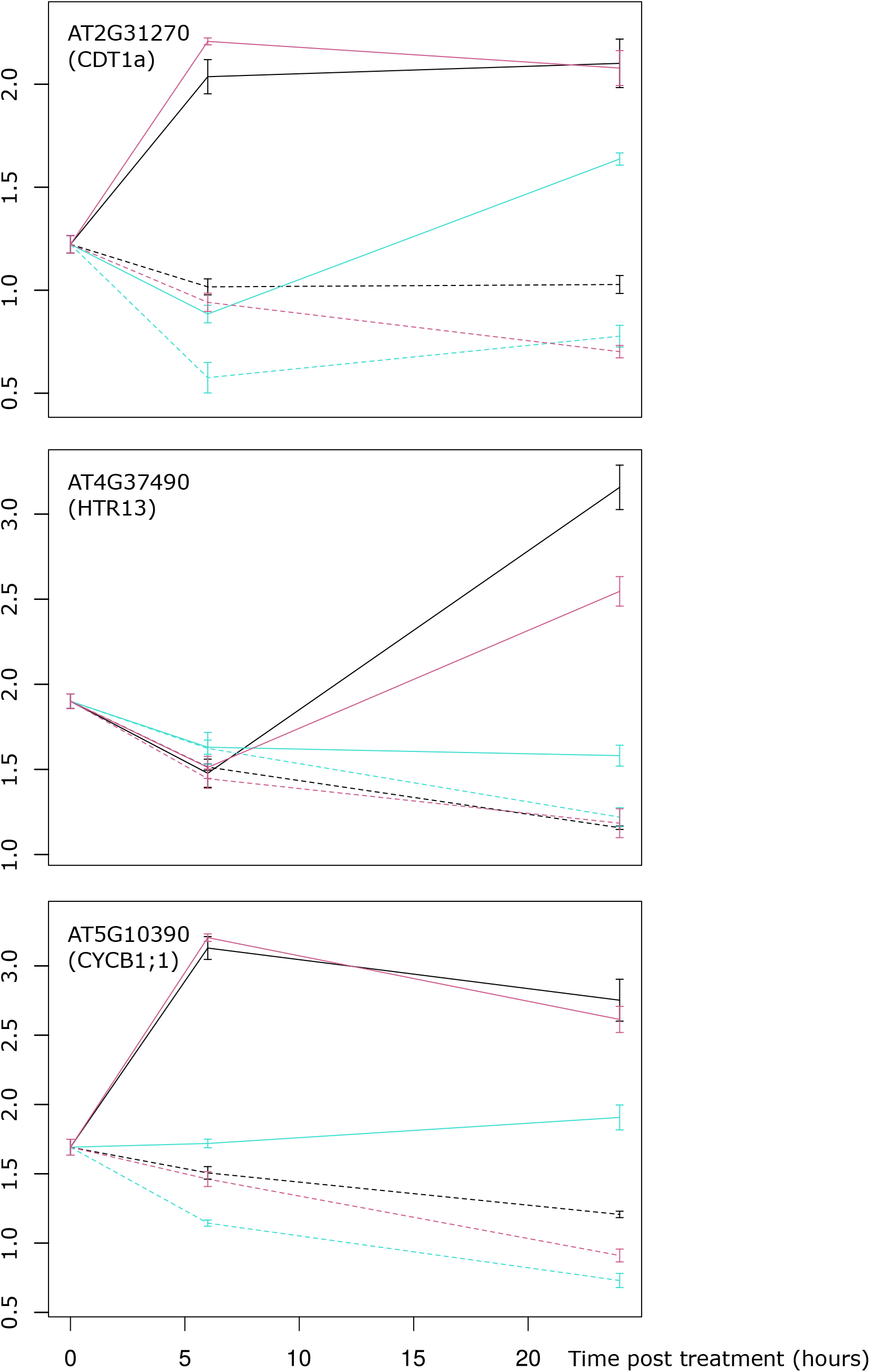
Expression profile of cell cycle marker genes (Desvoyes et al., 2020): CDT1a was shown to be specifically expressed in cells in G1, CycB1;1 was shown to be specifically expressed in cells in late G2 phase and early in mitosis while the histone HTR13 is expressed predominantly during S phase and early G2 phase.

**Supplemental Figure 10.**
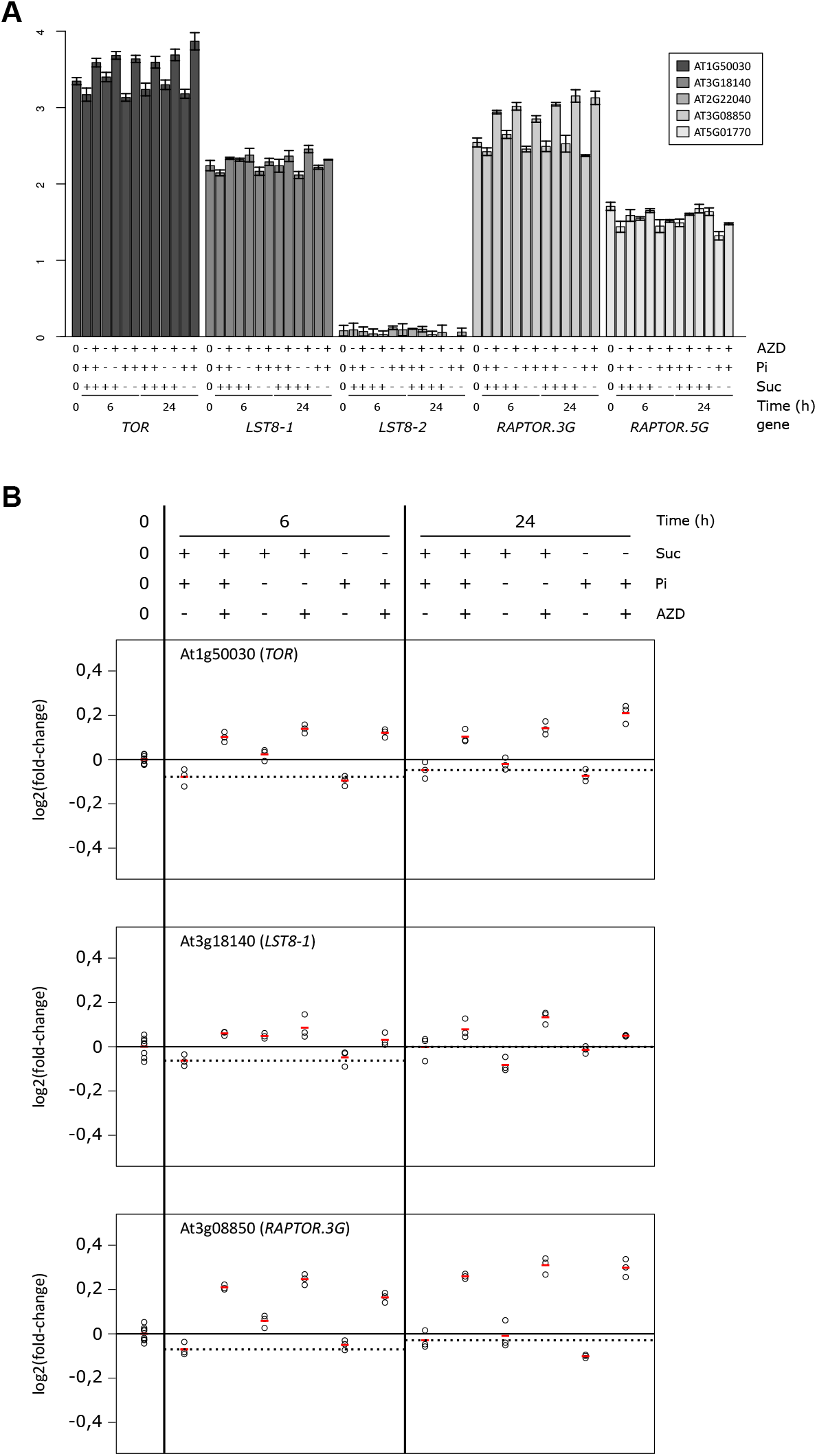
Expression profiles of TOR complex members. (A) Absolute expression of genes coding for components of the TOR complex. (B) Expression of genes coding for components of the TOR complex, relative to their expression values in the T0 samples. Circles represent individual values, red bars represent averages. The horizontal solid lines represent the T0 averages and the horizontal dashed lines represent averages of the corresponding non-treated samples.

**Supplemental Figure 11.**
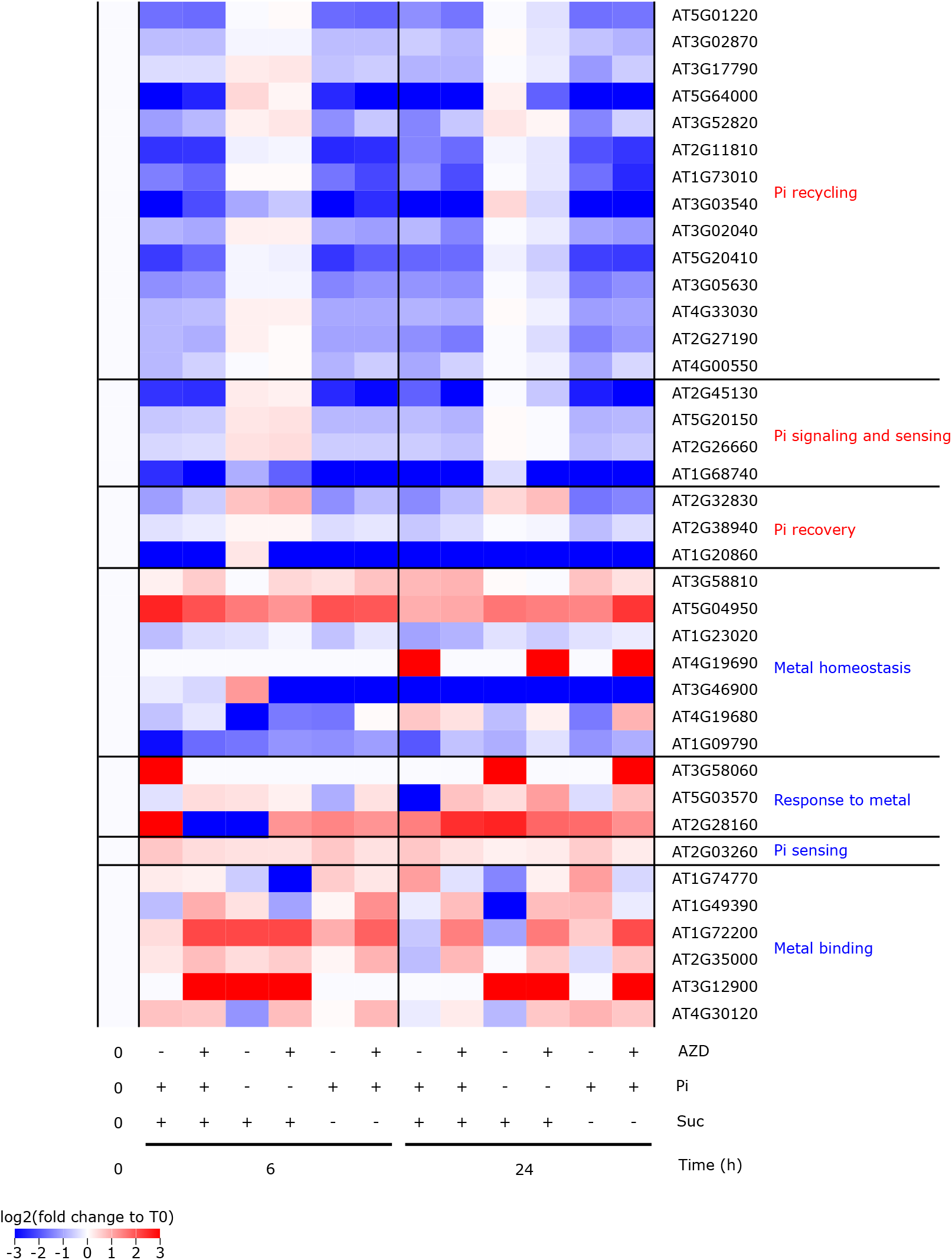
Expression profiles of phosphate starvation regulated genes. Expression profiles of the genes listed in Thibaud et al., (2010), normalized to their expression in the T0 samples. Annotations of genes on the right-hand side are from the same article (annotations in red and blue correspond to, repectively, genes described as upregulated after phosphate starvation and genes described as downregulated after phosphate starvation).

**Supplemental Figure 12.**
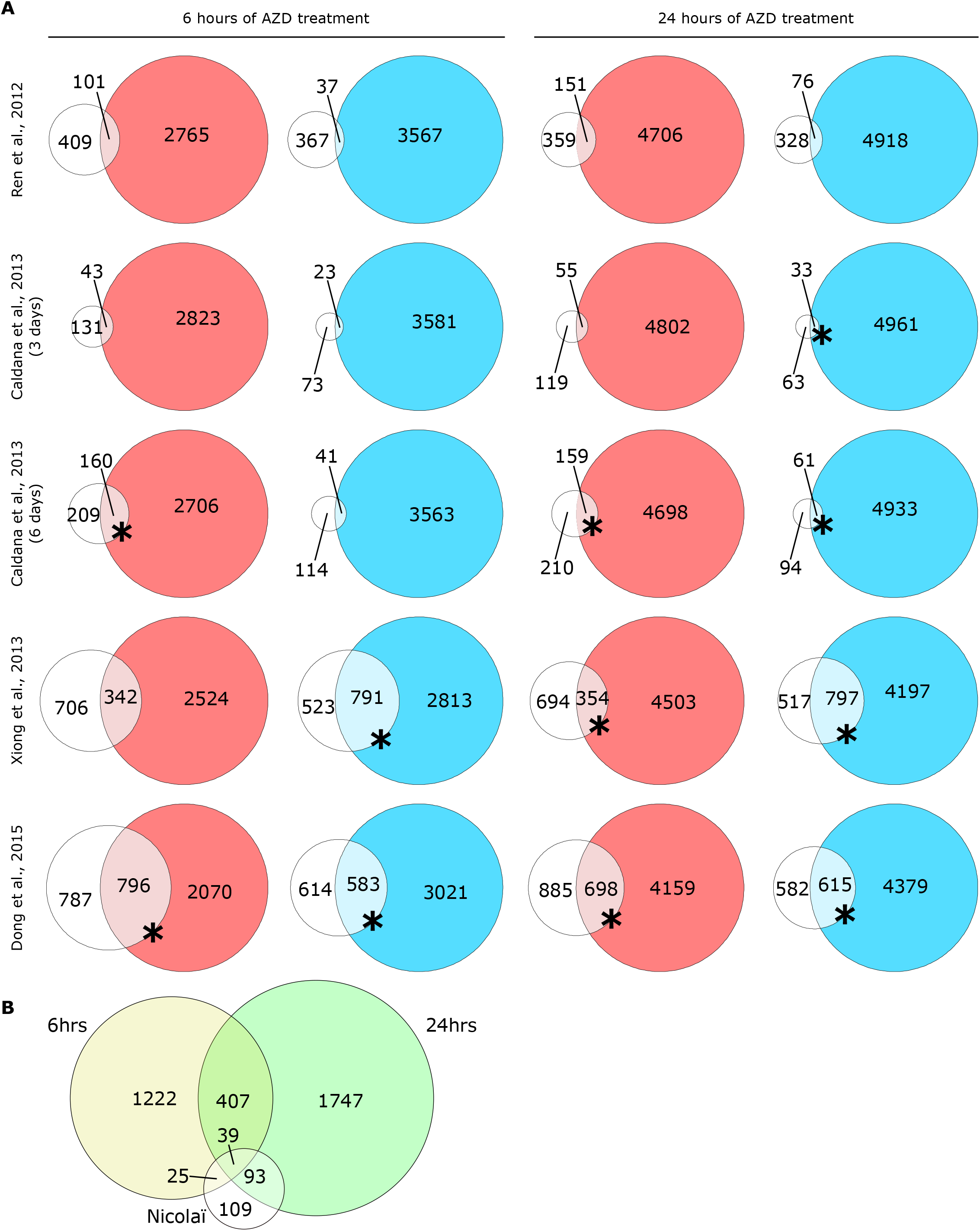
Comparison of our datasets with previously published datasets. (A) Comparison of genes found to be deregulated in response to AZD-8055 treatment in our experiment to the previously published data. Colored circles correspond to our dataset (red for upregulated genes and blue for downregulated genes). Stars represent datasets for which more than one third of the genes that they found to be deregulated are shared with our datasets. (B) Comparison of genes deregulated in response to 6 or 24 hours of sugar starvation in our experiment to the previously published work of Nicolaï et al. (2006)

Supplemental Material 1. Gene Ontology term enrichment of biological processes affected after 6 hours (A-B) or 24 hours (C-D) of nutrient resupply for the genes that are induced (A,C) or repressed (B,D) compared to the T0 condition. Only terms with an adjusted p-value ¡ 0.05 were kept for the analysis.

Supplemental Table 1. Medium composition for nutrient resupply (concentration given for the medium after dilution with the cells)

Supplemental Table 2. Metabolite changes during growth kinetics

Supplemental Table 3. Log2(fold change) in the evolution of metabolite content in 4-day old cells treated with 2 μM AZD-8055

Supplemental Table 4. Quantification of metabolites over time in response to medium re-supply (NPS), and phosphate (NS) and sucrose (NP) starvation, in the presence or absence of the TOR inhibitor AZD-8055 (AZD and DMSO respectively)

Supplemental Table 5. Quantification of lipids over time in response to medium re-supply (NPS), and phosphate (NS) and sucrose (NP) starvation in the presence or absence of the TOR inhibitor AZD-8055 (AZD and DMSO respectively)

Supplemental Table 6. List of deregulated gene expression for each of the pairwise comparisons. Log fold changes as well as adjusted p-values are displayed.

Supplemental Table 7. List of Gene Ontology terms enriched among the genes deregulated after nutrient re-supply compared to the T0 condition (only GO terms with an adjusted p-value ¦ 0.05 were kept). (A) Genes transcriptionally induced after 6 hours of nutrient resupply. (B) Genes transcriptionally repressed after 6 hours of nutrient resupply. (C) Genes transcriptionally induced after 24 hours of nutrient resupply. (D) Genes transcriptionally repressed after 24 hours of nutrient resupply. (xx corresponds to the number of GO occurrences in the dataset, X to the number of genes in the dataset, nn to the number of GO occurrences in the annotated genome, N to the size of the annotated genome)

Supplemental Table 8. List of Gene Ontology terms enriched among the genes deregulated between the different conditions compared by timepoint. (A) Genes upregulated after 6 hours of phosphate starvation. (B) Genes downregulated after 6 hours of phosphate starvation. (C) Genes upregulated after 6 hours of sugar starvation. (D) Genes downregulated after 6 hours of sugar starvation. (E) Genes upregulated after 6 hours of AZD-8055 treatment. (F) Genes downregulated after 6 hours of AZD-8055 treatment. (G) Genes upregulated after 6 hours of phosphate starvation and AZD-8055 treatment. (H) Genes downregulated after 6 hours of phosphate starvation and AZD-8055 treatment. (I) Genes upregulated after 6 hours of sugar starvation and AZD-8055 treatment. (J) Genes downregulated after 6 hours of sugar starvation and AZD-8055 treatment. (K) Genes upregulated after 24 hours of phosphate starvation. (L) Genes downregulated after 24 hours of phosphate starvation. (M) Genes upregulated after 24 hours of sugar starvation. (N) Genes downregulated after 24 hours of sugar starvation. (O) Genes upregulated after 24 hours of AZD-8055 treatment. (P) Genes downregulated after 24 hours of AZD-8055 treatment. (Q) Genes upregulated after 24 hours of phosphate starvation and AZD-8055 treatment. (R) Genes downregulated after 24 hours of phosphate starvation and AZD-8055 treatment. (S) Genes upregulated after 24 hours of sugar starvation and AZD-8055 treatment. (T) Genes downregulated after 24 hours of sugar starvation and AZD-8055 treatment. (xx corresponds to the number of GO occurrences in the dataset, X to the number of genes in the dataset, nn to the number of GO occurrences in the annotated genome, N to the size of the annotated genome)

Supplemental Table 9. List of Gene Ontology terms enriched among the treatment-specific genes deregulated after 24 hours of treatment when compared to the 24 hour control conditions. (A-G) Upregulated genes, (H-N) Downregulated genes. (A,H) specific to phosphate starvation, (B,I) common to phosphate starvation and AZD-8055 treatment but absent from sugar starvation treatment, (C,J) AZD-8055 treatment specific, (D,K) common to phosphate and sugar starvation but absent from the AZD-8055 treatment, (E,L) common to the 3 treatments, (F,M) common to the AZD-8055 treatment and sugar starvation but absent from phosphate starvation, (G,N) specific to sugar starvation. (xx corresponds to the number of GO occurrences in the dataset, X to the number of genes in the dataset, nn to the number of GO occurrences in the annotated genome, N to the size of the annotated genome)

## Footnotes

1. http://www.epigenesys.eu/en/protocols/bio-informatics/1283-guidelines-for-rna-seq-data-analysis

2. http://www.bioinformatics.babraham.ac.uk/projects/fastqc/

3. Anders S. 2010. HTSeq: Analysing high-throughput sequencing data with python. URL https://htseq.readthedocs.io/en/master/

4. https://www.ebi.ac.uk/ena/browser/home

5. https://dx.doi.org/10.5281/zenodo.4608028

